# A Multi-Kinase Inhibitor Screen Identifies mTOR Signaling as a Vulnerability in Advanced Prostate Cancer

**DOI:** 10.64898/2026.10.13.758818

**Authors:** Sriya Sridhar, Kabhilan Mohan, Smitha Kizhake, Amarnath Natarajan, David L Kelly, Grinu Mathew

**Affiliations:** Eppley Institute for Research in Cancer and Allied Diseases, University of Nebraska Medical Center, Omaha, NE, 68198, USA; Division of Pharmaceutical Sciences, James L Winkle, College of Pharmacy, University of Cincinnati, Cincinnati, OH 45267 USA; Department of Pharmaceutical Sciences, University of Nebraska Medical Center, Omaha, NE 68198 USA; Department of Genetics, Cell Biology and Anatomy, University of Nebraska Medical Center, Omaha, NE 68198 USA; Nebraska Center for Molecular Target Discovery and Development, University of Nebraska Medical Center, Omaha, NE, USA; Fred and Pamela Buffett Cancer Center, University of Nebraska Medical Center, Omaha, NE, USA

**Keywords:** Receptor tyrosine kinase AXL, prostate cancer, kinase inhibitor screen, PI3K/AKT/mTOR, Torin-2, PRISM DepMap, drug repurposing

## Abstract

The receptor tyrosine kinase AXL has context-dependent roles in prostate cancer (PCa). Although AXL signaling has been implicated in PCa progression, it can also promote disseminated tumor cell dormancy. Paradoxically, *AXL* expression has been reported to be lost in metastatic PCa. This raises two critical questions: Is AXL loss associated with aggressive disease, and does it create therapeutically exploitable vulnerabilities? In this study, we comprehensively analyzed *AXL* expression across pan-cancer and PCa cohorts and found that *AXL* downregulation was significantly associated with treatment resistance, aggressive PCa phenotypes, and shorter overall survival. To investigate the functional consequences of AXL loss, we used CRISPR-Cas9 based gene editing to generate isogenic *AXL*^KO^ cells and characterized their proliferative and migratory properties. To identify therapeutically exploitable vulnerabilities associated with AXL loss, we screened these cells against a library of 1,814 kinase inhibitors using the isogenic cell lines. We identified Torin-2 as a candidate therapeutic agent and validated its activity using dose-response assays, which demonstrated an ∼ 18-fold lower IC_50_ in *AXL*^KO^ cells than in *AXL*^WT^ cells. In parallel, PRISM/DepMap analysis provided orthogonal evidence of drug dependencies associated with low AXL expression and revealed a selective vulnerability of *AXL*^KO^ cells to PI3K/AKT/mTOR pathway inhibition. Collectively, our findings link AXL downregulation to aggressive PCa phenotypes and identify mTOR inhibition as a potential therapeutic strategy for AXL-deficient PCa.

**Significance:** AXL downregulation is associated with aggressive PCa, treatment resistance, and shorter overall survival, highlighting its potential clinical significance. By identifying a selective vulnerability of AXL-deficient cells to mTOR inhibition, this study reveals a potential therapeutic strategy for AXL-low PCa.

Graphical abstract

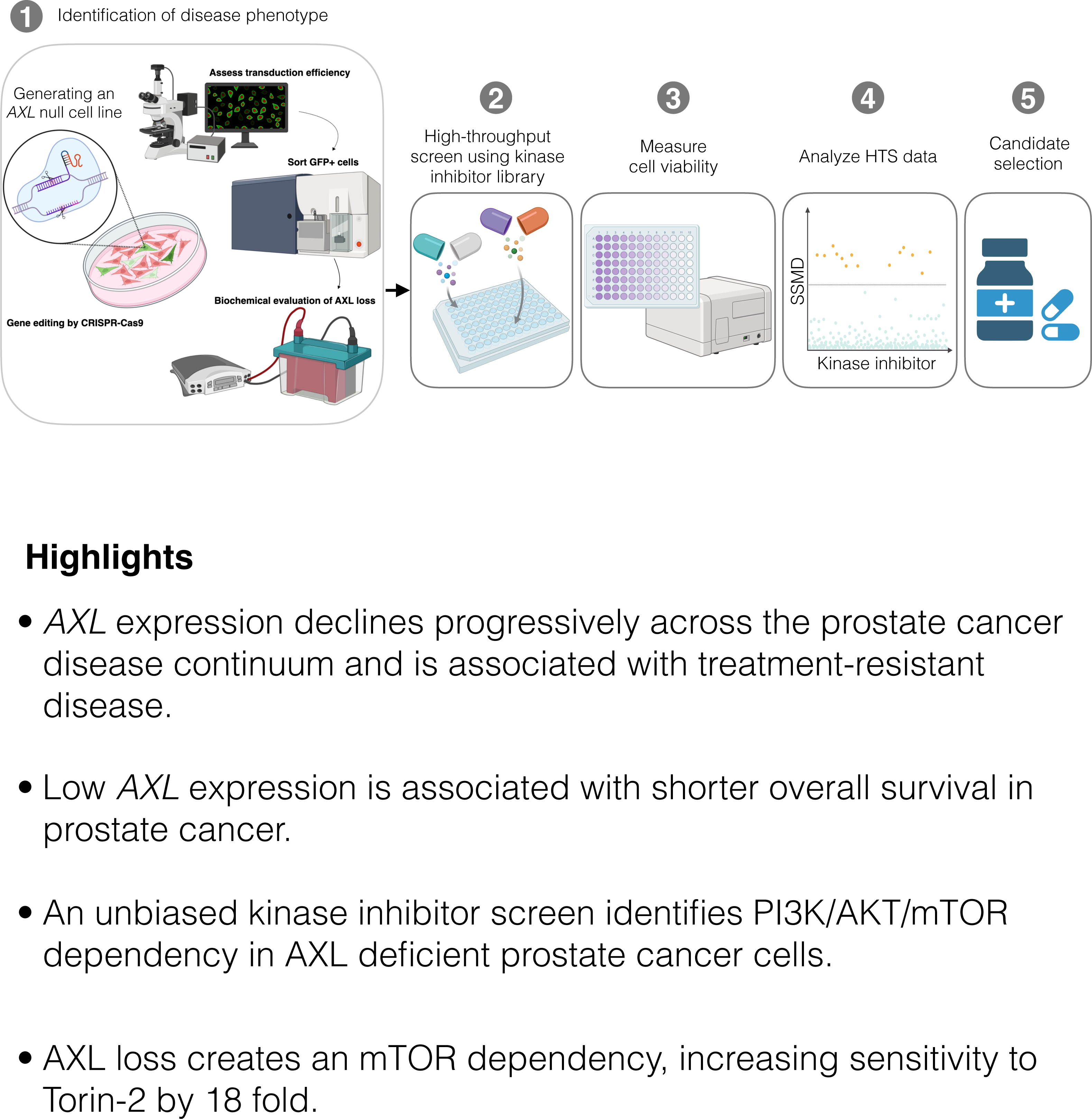

## Introduction

Metastatic prostate cancer (PCa) remains largely incurable, with mortality driven by the emergence and progression of disseminated disease [1, 2]. A defining feature of PCa metastasis is the ability of disseminated tumor cells (DTCs) to persist in distant tissues in a quiescent state before re-entering the cell cycle and giving rise to overt metastatic lesions [3–5]. The transition from cellular quiescence to active proliferation is therefore a critical step in metastatic progression. Identifying signaling dependencies that emerge when dormant cells exit dormancy and resume proliferation may reveal new therapeutic opportunities to prevent metastatic disease.

In PCa, AXL, a receptor tyrosine kinase in the TAM (TYRO3, AXL, MERTK) family [6–8], has emerged as an important regulator of the balance between dormancy and proliferative outgrowth [9–13]. AXL is activated primarily by the vitamin K-dependent ligand growth arrest-specific 6 (GAS6) [14] and regulates cellular survival, proliferation, migration, and adaptation to the tissue microenvironment [15, 16]. Although AXL is widely regarded as a pro-tumorigenic receptor that promotes invasion, metastasis, immune evasion, and therapeutic resistance across multiple malignancies [17–26], studies in PCa have revealed a distinct and context-dependent function. For example, within the bone marrow niche, osteoblast-derived GAS6 binds to AXL on DTCs to suppress proliferation and promote a quiescent state [9]. AXL is also required for TGF-β2-mediated dormancy induction, whereas MERTK can promote dormancy escape through MAP kinase signaling [12, 27, 28]. Consistent with these findings, the relative balance of AXL and TYRO3 has also been shown to function as a molecular switch governing DTC fate; cells with higher AXL expression remain largely quiescent [10]. Furthermore, Axelrod et al. demonstrated that AXL is necessary for PCa cells to enter dormancy, with AXL overexpression reducing EdU incorporation, restricting cell proliferation, and delaying tumor formation *in vivo* [13]. Notably, they showed that AXL is largely absent from tumor cells in human primary and metastatic PCa and that higher AXL expression is associated with better overall survival [13].

Dormant DTCs are difficult to eliminate because of their low proliferative activity and relative resistance to conventional cytotoxic therapies [29–31]. Conversely, escape from dormancy enables proliferative outgrowth and the development of clinically overt metastases, which remain incurable in PCa [32–34]. These observations raise an important therapeutic question: how can cells that escape AXL associated dormancy be targeted? Because tumor-cell AXL may already be lost in advanced PCa [13], strategies designed to inhibit AXL activity may have limited relevance in this setting. We reasoned instead that AXL loss may represent a transition to a distinct cellular state, accompanied by adaptive signaling changes that support continued proliferation and metastatic progression. Loss of a receptor tyrosine kinase can alter signaling across interconnected kinase networks, enabling alternative receptors and downstream pathways to compensate for the missing signal [18, 35]. Although such adaptation may facilitate tumor progression, it may also create selective signaling dependencies that can be therapeutically exploited.

In this study, we show that AXL expression declines during PCa progression and is associated with aggressive metastatic disease. To identify dependencies in AXL-deficient PCa, we performed a systematic kinase inhibitor screen using isogenic AXL-proficient and AXL-deficient PCa cells. Our approach revealed preferential sensitivity of AXL-deficient cells to inhibition of the PI3K/AKT/mTOR signaling network. Pharmacologic validation of the screen hits identified the ATP competitive mTOR inhibitor Torin-2 as the most potent genotype-selective compound. Orthogonal analysis of the DepMaP PRISM repurposing dataset [36–38] further supported a dependence on the PI3K/AKT/mTOR signaling axis. Together, these findings establish *AXL* loss as a distinct signaling state associated with aggressive PCa and identify a therapeutically actionable dependency that emerges following AXL loss.

## Materials and Methods

### Cell Culture

PC3 cells were maintained in RPMI medium supplemented with 10% fetal bovine serum (FBS), 100 µg/mL penicillin, 100 µg/mL streptomycin, and 2 mM L-glutamine. Cell lines were cultured as 2D monolayers in cell culture-grade Petri dishes and maintained at 37°C in a humidified incubator with 5% CO_2_. Routine checks for potential mycoplasma infection were conducted using the LookOut One-Step Mycoplasma PCR Detection Kit (Cat# MP0050-100TST) from Sigma-Aldrich.

### Generation and validation of AXL knockout cells

*AXL*^KO^ PC3 cells were generated using CRISPR-Cas9 gene editing. Three guide RNAs (gRNAs) targeting exons 2, 8, and 20 of human *AXL* were used: exon 2, 5′-AACCTGGAGCTGACACCGAA-3′ [13]; exon 8, 5′-CGCGTAGCACTAATGTTCTC-3′; and exon 11, 5′-ATGCCTTGATGTCGCGGTGC-3′. The gRNAs were cloned into the LRG2.1 sgRNA vector (Addgene #108098) using the CRISPR cloning strategy described by Tarumoto et al. [39]. PC3 cells were transfected with the *AXL*-targeting LRG2.1 plasmids together with a Cas9-GFP expression plasmid (Addgene #52962). GFP-positive cells were isolated by fluorescence-activated cell sorting, seeded at low density, and expanded as individual clones. Clones were screened by sequencing of the *AXL* target loci to identify CRISPR-induced indels. A control sg*ROSA26* clone was generated using the corresponding control CRISPR plasmid (Addgene #105587). All plasmids were obtained from Addgene.

### Flow Cytometry

Cells were detached using trypsin-EDTA and washed in FACS buffer [PBS containing 2% FBS/BSA], and stained with a PE-conjugated anti-human AXL antibody (R&D FAB154P) or PE-conjugated isotype-matched IgG control (R&D IC002P) for 1 hour at 4 °C in the dark. Cells were washed twice and analyzed using a BD LSRFortessa flow cytometer. PE fluorescence was detected using the 561-nm laser and an appropriate yellow-green detector. At least 100,000 events were collected per sample following exclusion of debris and doublets based on FSC/SSC. Data were analyzed using FlowJo. PE-positive and PE-negative gates were established using the isotype control and applied identically to *AXL*^WT^ and *AXL*^KO^ cells.

### Immunoblotting

Whole cell lysates were prepared in RIPA buffer supplemented with protease and phosphatase inhibitors. Protein concentrations were determined using a BCA protein assay, and 15 µg of protein was separated by SDS-PAGE and transferred to PVDF membranes. Membranes were blocked with 5% BSA in TBST and incubated overnight at 4 °C with antibodies against AXL (Cell Signaling Technology,Cat# 8661, 1:1000) and β-actin (Sigma-Aldrich, A3854, 1:15,000). Following incubation with HRP-conjugated secondary antibodies, protein bands were detected by enhanced chemiluminescence using a ChemiDoc Imaging System (Bio-Rad). Lysates from the immortalized normal human prostate epithelial cell line PNT2-C2 were included as an AXL-positive reference control.

### Proliferation assay

*AXL*^WT^ and *AXL*^KO^ cells were seeded at 5000 cells per well in 96-well plates containing RPMI medium with 10% FBS, with 4 replicate wells per condition. Cell growth was monitored every 6 h for 72 h using a Cytation 10 Cell Imaging Multimode Reader (Agilent BioTek) equipped with a BioSpa 8 Automated Incubator and maintained at 37 °C with 5% CO₂. Brightfield images were acquired using a 4X objective. Cell numbers were quantified using the Cellular Analysis module in Gen5 software with object size/intensity thresholds. Cell counts were normalized to the maximum cell count across all conditions and expressed as a percentage of the maximum cell number. Percent cell count between *AXL*^WT^ and *AXL*^KO^ cell lines at each time point was analyzed using two-way repeated-measures ANOVA with the Geisser-Greenhouse correction, followed by Sidak’s multiple comparisons test. A *P* value below 0.05 was considered statistically significant. Statistical significance was defined * *P* < 0.05, ** *P* < 0.01, *** *P* < 0.001, and **** *P* < 0.0001.

### Migration assay

*AXL*^WT^ and *AXL*^KO^ cells were seeded at 10,000 cells per well in 96-well plates and grown to confluence. Prior to the scratch assay, cells were treated with Mitomycin C (Sigma-Aldrich, 10 µg/ml for 2hrs) to inhibit cell proliferation, ensuring that the wound closure data reflected cell migration. Uniform wounds were generated using the Agilent BioTek AutoScratch Wound Making Tool. Wells were washed twice with PBS to remove detached cells, and fresh medium was added. Plates were imaged immediately after wounding (0 h) and every 6 h for 48 h using the Cytation 10 Cell Imaging Multimode Reader (Agilent BioTek) under the same environmental conditions. Wound closure was quantified using Gen5 software, and migration was expressed as the percentage of wound area closed relative to the initial wound area. Wound closure data in *AXL*^WT^ and *AXL*^KO^ cell lines was analyzed using two-way repeated-measures ANOVA with the Geisser-Greenhouse correction, followed by Sidak’s multiple comparisons test. A *P* value below 0.05 was considered statistically significant. Statistical significance was defined * *P* < 0.05, ** *P* < 0.01, *** *P* < 0.001, and **** *P* < 0.0001.

### Transcriptomic analysis of AXL expression across pan-cancer cohorts and prostate cancer disease states

*AXL* mRNA expression was obtained from recount3 [40], which provides uniformly reprocessed TCGA RNA-seq data generated through the Monorail pipeline using STAR alignment to GRCh38 and GENCODE v26 gene annotation. For each of the 33 TCGA projects, project-level gene-count matrices and corresponding sample metadata were downloaded from the recount3 repository. Tissue type was assigned using the GDC-derived metadata field “gdc_cases.samples.sample_type”, which classifies samples as Solid Tissue Normal, Primary Tumor, or Metastatic. The complete dataset comprised of 11,348 samples across 33 cancer types, including 9,962 primary tumor, 742 solid-tissue-normal, and 394 metastatic samples. *AXL* gene level read counts were normalized within each sample to counts per million (CPM) by dividing the *AXL* count by the total gene-level read sum for that sample and multiplying by 10^6^. Values were then log_2_ transformed as log_2_ CPM. Analyses were restricted to the 16 TCGA cancer types containing at least 10 solid-tissue-normal samples: ACC, DLBC, GBM, LGG, MESO, OV, PAAD, PCPG, SARC, SKCM, TGCT, THYM, UCS, UVM, LAML, and CESC were excluded because of insufficient normal tissue. Cancer types were ordered according to decreasing median *AXL* expression in primary tumors. Two complementary analyses were performed using the same normalized expression data. Patient-matched analysis was restricted to patients contributing both a solid-tissue-normal and primary-tumor sample, allowing each patient to serve as their own reference. Patient identity was defined using the first 12 characters of the TCGA barcode. When multiple aliquots from the same patient and tissue type were available, *AXL* expression values were averaged to generate a single patient-level value before pairing. This yielded 692 matched normal-tumor pairs across the 16 cancer types. Within each cancer type, primary-tumor and matched normal *AXL* expression was compared using a two-sided Wilcoxon signed-rank test, with the paired effect size calculated as the median within-patient difference (primary tumor-normal). We also compared the complete solid-tissue-normal and primary-tumor cohorts within each cancer type without requiring patient matching. This analysis included 713 solid-tissue-normal and 7,189 primary-tumor samples across the 16 cancer types. *AXL* expression was compared between the two groups using a two-sided Mann-Whitney U test, with the effect size calculated as the difference in group medians (primary tumor-normal). For both analyses, box plots show the median and interquartile range (IQR), with whiskers extending to the most extreme value within 1.5 x IQR; outliers were not displayed. For each analysis, the 16 cancer-type-specific *P* values were adjusted for multiple comparisons using the Benjamini-Hochberg false-discovery-rate (FDR) procedure. Statistical significance was defined as FDR < 0.05, with FDR < 0.05, <0.01, and <0.001 indicated by *, **, and ***, respectively.

Transcriptomic analysis of AXL, MYC, EZH2, and MKI67 across PC disease states. RNA-sequencing data was obtained from the NCI Genomic Data Commons (GDC) Data Portal (https://portal.gdc.cancer.gov). Cases were selected using the following criteria: primary site = prostate gland, tissue or organ of origin = prostate gland, and data access = open. The resulting cohort included 691 RNA-seq samples from 634 patients, comprising samples from The Cancer Genome Atlas Prostate Adenocarcinoma project (TCGA-PRAD) and the West Coast Prostate Cancer Dream Team metastatic castration-resistant prostate cancer project (WCDT-MCRPC; dbGaP phs001648). Samples were classified according to GDC sample type as solid-tissue normal (n = 52), primary tumor (n = 539), or metastasis (n = 100). Gene-level raw counts were obtained from GDC STAR-Counts files, generated by alignment to GRCh38 using STAR and quantification against the GENCODE gene annotation. Count matrices were merged and analyzed using DESeq2. Samples were normalized using the DESeq2 variance-stabilizing transformation (VST; blind = FALSE), with sample type (normal, primary tumor, or metastasis) and study of origin included in the DESeq2 design formula (∼batch + sample_type). VST-transformed expression values for *AXL, MYC, EZH2,* and *MKI67* were used for visualization and downstream comparisons. Expression was compared between normal tissue and primary tumors and between primary tumors and metastatic samples using two-sided Wilcoxon rank-sum tests. *P* values were adjusted using the Benjamini-Hochberg false-discovery-rate procedure. Violin plots show the distribution of VST-transformed expression values; individual points represent samples, and horizontal bars indicate group medians.

Expression of *AXL* in relation to clinical and molecular features was obtained from the Prostate Cancer Atlas (PCaProfiler; https://www.prostatecanceratlas.org) [41]. The atlas is a compendium of publicly available bulk RNA-sequencing data from normal prostate, primary tumors, and metastatic castration-resistant prostate cancer. We used transcript-level expression of the *AXL* transcript (Ensembl/GENCODE). Raw sequencing data from the contributing studies were uniformly processed against GRCh38 using the atlas pipeline, and transcript-level expression was normalized and harmonized across datasets. Normalized log_2_ CPM expression values and corresponding clinical and molecular annotations were obtained from the atlas for downstream analysis. Statistical analyses for the PCaProfiler cohort were performed according to the clinical or molecular feature examined, using the statistical tests specified for each comparison. Two-group comparisons were performed using two-sided Wilcoxon rank-sum tests against the indicated reference groups. For multi-group comparisons, overall differences were assessed using the Kruskal-Wallis test, followed by pairwise Wilcoxon rank-sum tests comparing each disease stage or molecular subtype with normal prostate tissue. *P* values were adjusted using the Benjamini-Hochberg procedure. Samples lacking relevant clinical annotations were excluded from the corresponding analysis. A *P* value below 0.05 was considered statistically significant. Statistical significance was defined * *P* < 0.05, ** *P* < 0.01, *** *P* < 0.001, and **** *P* < 0.0001.

*AXL* and *GAS6* mRNA expression across nine PCa cell lines was obtained from a previously published RNA-seq dataset by Smith et al. [42]. The panel included AR-indifferent (DU145, PC3, PC3M), AR-responsive (VCaP, CWR22Rv1, 22Rv1, LNCaP, and LNCaP-C4-2B), and neuroendocrine (NCI-H660) cell lines. Expression values were visualized as a heatmap using GraphPad Prism.

### Survival Analysis

Overall survival was analyzed using the survival module of the Prostate Cancer Atlas. Patients with available follow-up data (n = 661) were dichotomized according to median AXL transcript expression (log_2_ CPM = 4.73). Survival probabilities were estimated using the Kaplan–Meier method, and hazard ratios (HRs) with 95% confidence intervals (CIs) for high versus low AXL expression were estimated using Cox proportional hazards regression. *P* values were calculated using the log-rank test. Survival differences between patients with and without *AXL* genomic alterations were assessed using the log-rank test in cBioPortal.

### Kinase inhibitor screen and hit identification

PC3 *AXL*^WT^ and PC3 *AXL*^KO^ cells were independently seeded in 96 well plates using an automated liquid dispenser to ensure uniform cell seeding. After 24 h, cells were treated with compounds from the SelleckChem kinase inhibitor library containing 1,930 compounds distributed across 22 plates (Cat#L1200). Up to 88 compounds were tested per plate at final concentrations of 0.5 or 2.5 µM. Each cell line and drug concentration was tested in three independent replicate plates. Each screening plate included bemcentinib (BGB324/R428; SelleckChem) at 10 µM as a positive control for cytotoxicity (column 12, rows C-D), alongside vehicle (DMSO or water) negative-control wells. Bemcentinib is a selective AXL inhibitor at nanomolar concentrations. At 10 µM it was used only as a plate-level positive control for loss of viability, because at this concentration it is cytotoxic through off-target activity, independent of AXL. Consistent with this, it reduced viability to a similar extent in PC3 *AXL*^WT^ and PC3 *AXL*^KO^ cells. Positive-control wells were used to confirm the assay window on each plate and were not used to calculate SSMD. The vehicle control consisted of the highest concentration of DMSO used for compounds on the corresponding plate. Cells were exposed to compounds for 72 h, after which cell viability was assessed by MTT assay. Drug effects were quantified using the strictly standardized mean difference (SSMD), as previously described [43]. SSMD values were calculated as: SSMD = (mean drug - mean vehicle) / √(SD_drug^2^ + SD_vehicle^2^). Compounds with SSMD < −3 were considered to have strong inhibitory effects on cell viability. Compounds were classified as affecting both cell lines if they met the SSMD threshold in all three replicate plates at both concentrations in both PC3 sgROSA and PC3 *AXL*^KO^ cells. Compounds meeting the threshold in only one cell line at both concentrations were classified as cell-line selective hits.

Drug targets were assigned based on the target annotations provided by SelleckChem. Compounds in the kinase inhibitor library were classified into 20 primary target pathways plus “Others” using vendor annotations. *AXL*^KO^ selective hits were defined as compounds meeting the hit threshold (SSMD < −3.0) in *AXL*^KO^ but not *AXL*^WT^ cells. Pathway-specific hit rates were calculated as the percentage of annotated compounds identified as selective hits and compared with the screen-wide background hit rate (1.5%). Pathway enrichment was assessed using a one-sided hypergeometric test, with P values adjusted using the Benjamini-Hochberg procedure (FDR < 0.05).

### PRISM drug sensitivity analysis and integration with the kinase inhibitor screen

The dose-response data from the PRISM repurposing screen (20Q2) were obtained from the DepMap portal [38], and gene expression data were obtained from the DepMap Public 24Q4 dataset. Cell lines represented in the PRISM dataset were stratified by *AXL* mRNA expression into *AXL*-Low (first quartile; log_2_ (TPM + 1) < 0.4; n = 40) and *AXL*-High (fourth quartile; log_2_ (TPM + 1) > 6.4; n = 155) groups. For each compound, area under the dose-response curve (AUC) values were converted to drug-specific z-scores across cell lines, and cell lines with z-scores < −1 were classified as sensitive. An unbiased analysis was first performed across all 1,448 compounds in the PRISM dataset. For each compound, the proportions of sensitive cell lines in the *AXL*-Low and *AXL*-High groups were compared using two-sided Fisher’s exact tests to identify associations between *AXL* expression and drug sensitivity independently of the kinase inhibitor screen. The PRISM compound list was then intersected with the hits identified in the kinase inhibitor screen. Seven overlapping compounds, bardoxolone methyl, taselisib, dasatinib, Torin-2, pictilisib (GDC-0941), VE-822, and MK-8745, were selected for comparative analysis of drug sensitivity between *AXL*-Low and *AXL*-High cell lines. *P* values were reported without adjustment for multiple comparisons.

### Dose-response analysis of hit compounds

PC3 *AXL*^WT^ or *AXL*^KO^ cells were seeded in 96-well plates at 10,000 cells per well and allowed to adhere overnight. Cells were then treated with compounds using 2-fold serial dilutions from compound-specific top concentrations. For bardoxolone methyl, cinobufagin, dasatinib, MCB-613, nilotinib, SGC-GAK-1, SR-4835, tomatine, and Torin-2, we tested 12 concentrations across 11 successive 2-fold dilutions, plus a 0.1 nM anchor point. Top concentrations were 2.5 µM for cinobufagin, dasatinib, tomatine, and Torin-2; 5 µM for MCB-613, nilotinib, SGC-GAK-1, and SR-4835; and 10 µM for bardoxolone methyl. For taselisib, uprosertib, and VE-822, we tested nine concentrations across eight successive 2-fold dilutions from a top concentration of 30 µM, plus a 0.1 nM anchor point. Vehicle-control wells received DMSO at a concentration equivalent to the highest compound concentration tested. Each concentration was tested in technical triplicate across 3-4 independent biological replicates. After 72 h of compound exposure, cell viability was assessed by MTT assay. Raw optical density values were normalized to the mean vehicle-control signal on the corresponding plate and expressed as percentage cell viability. Dose-response curves were fitted in GraphPad Prism using a variable-slope nonlinear regression model (log [inhibitor] versus normalized response), with concentrations entered on a log_10_ scale. IC_50_ values and their 95% confidence intervals were obtained from the fitted curves. Maximal growth inhibition was assessed using the observed cell viability at the highest concentration tested. Genotype selectivity was quantified as the IC_50_ fold-shift (*AXL*^WT^ IC_50_/ *AXL*^KO^ IC_50_), with values > 1 indicating greater sensitivity of *AXL*^KO^ cells. Fold-shift 95% confidence intervals and *P* values were derived from the difference in log_10_(IC_50_) between genotypes using a two-sided Wald test. *P* values were adjusted across the 11 evaluable compounds using the Benjamini-Hochberg procedure. Compounds with an IC_50_ fold-shift ≥ 1.3 and an adjusted *P* < 0.05 were classified as selective for AXL loss. Nilotinib was excluded from this analysis because its IC_50_ exceeded the highest concentration tested in both genotypes. Data are presented as mean ± SD.

### Software and statistical analysis

The number of independent experimental replicates is indicated in the corresponding figure legends or main text. Statistical analyses and data processing was performed in R package and GraphPad PRISM. Data analysis in Python 3.13 using pandas, NumPy, SciPy, and statsmodels was performed using Claude (Anthropic, San Francisco, CA), an AI-assisted research tool. Box plots were generated using Matplotlib and exported as 600-dpi LZW-compressed TIFF files.

## Results

### AXL Expression is Broadly Dysregulated Across Human Cancers and Declines with Prostate Cancer Progression

To determine whether AXL expression is altered in cancer, we compared AXL mRNA levels in solid tissue “normal” and primary-tumor samples (Fig. 1A) across TCGA cancer types profiled by recount3 [40]. Sixteen cancer types had sufficient matched normal tissue (n ≥ 10 normal samples; Supplemental Fig. 1A), enabling a paired-cohort comparative analysis comprising 713 normal and 7,189 primary-tumor samples. *AXL* expression was significantly upregulated in tumors relative to normal tissue in kidney renal clear cell carcinoma (KIRC; n = 72 normal and n = 543 tumor), kidney renal papillary cell carcinoma (KIRP; n = 32 normal and n = 290 tumor), thyroid carcinoma (THCA; n = 59 normal and n = 505 tumor), and head and neck squamous cell carcinoma (HNSC; n = 44 normal and n = 502 tumor) (all *P* < 0.001). Conversely, *AXL* expression was significantly downregulated in the majority of tumor types examined, including breast (BRCA; n = 114 normal and n = 1127 tumor), lung adenocarcinoma (LUAD; n = 59 normal and n = 540 tumor), lung squamous cell carcinoma (LUSC; n = 51 normal and n = 504 tumor), prostate adenocarcinoma (PRAD; n= 52 normal and n = 505 tumor; bold in red), bladder (BLCA; n = 19 normal and n = 414 tumor), colon (COAD; n = 41 normal and n = 503 tumor), rectal (READ; n = 10 normal and n = 166 tumor), uterine corpus endometrial (UCEC; n = 35 normal and n = 553 tumor), kidney chromophobe (KICH; n = 25 normal and n = 66 tumor), and liver hepatocellular carcinoma (LIHC; n = 50 normal and n = 371 tumor) (all *P* < 0.001). Stomach (STAD; n = 37 normal and n = 416 tumor) and esophageal (ESCA; n = 13 normal and n = 184 tumor) carcinomas showed no significant difference. Taken together, our pan-cancer analysis demonstrates that *AXL* expression is not uniformly upregulated across human cancers at the transcriptional level. Instead, its expression is highly tissue context-dependent, with consistent upregulation in renal and head and neck carcinomas but significant downregulation in several epithelial malignancies, including endometrial, bladder, lung, and PCa.

**Figure 1.**
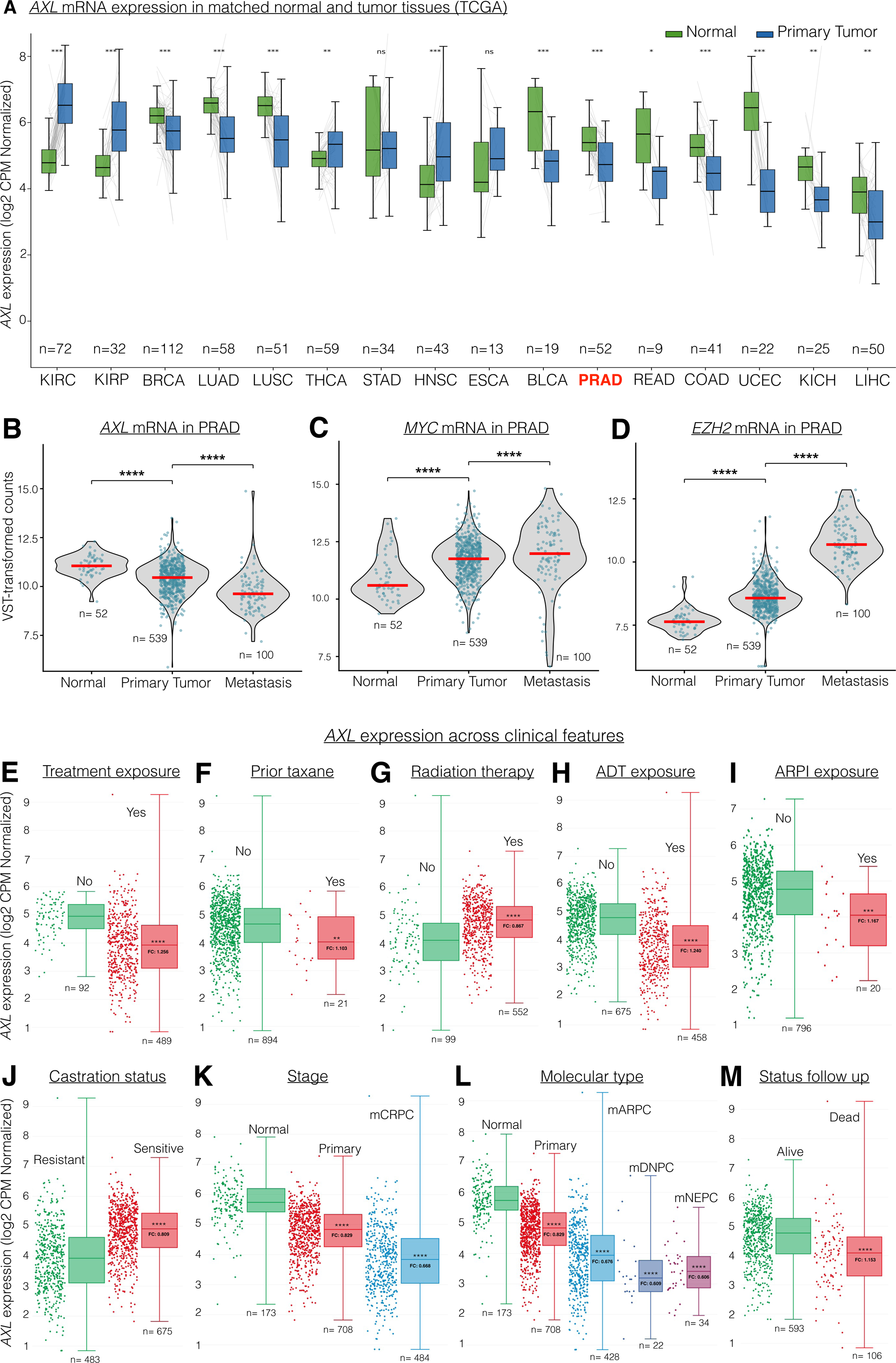
AXL mRNA expression is progressively lost across prostate cancer progression, treatment exposure, and AR-independent lineage transition. **(A)** AXL mRNA expression in matched normal and primary tumor tissues across 16 TCGA cohorts, shown as log2 (CPM+1) normalized counts. Cohorts are ordered from left to right by decreasing median tumor expression. Sample sizes are indicated as *n* (normal, tumor): KIRC (72), KIRP (32), BRCA (112), LUAD (58), STAD (34), THCA (59), HNSC (43), LUSC (51), ESCA (13), PRAD (52; bold red), BLCA (19), COAD (41), READ (9), UCEC (22), KICH (25), and LIHC (50). Boxes indicate the interquartile range with the median shown as a horizontal line; whiskers extend to 1.5 x IQR. AXL is significantly elevated in KIRC, KIRP, THCA, and HNSC, and significantly reduced in BRCA, LUAD, LUSC, PRAD, BLCA, COAD, READ, UCEC, KICH, and LIHC. **(B-D)** *AXL*, *MYC*, and *EZH2* mRNA expression across PCa disease progression, shown as VST-transformed counts. Groups include normal prostate (n = 52), primary tumor (n = 539), and metastasis (n = 100). Individual samples are shown as points, violin plots depict the distribution, and red horizontal bars indicate group medians. Pairwise comparisons are indicated by brackets. *AXL* decreases across progression, whereas *MYC* and *EZH2* increase. *AXL* mRNA expression in an expanded prostate adenocarcinoma cohort stratified by clinical and molecular annotations. Individual samples are shown as jittered points with box-and-whisker summaries; boxes indicate the interquartile range, horizontal lines indicate medians, and whiskers extend to the data range. **(E)** Treatment-naive (n = 92) versus previously treated (n = 489). **(F)** No prior taxane exposure (n = 894) versus taxane exposure (n = 21). **(G)** No radiation exposure (n = 99) versus radiation exposure (n = 552). **(H)** No ADT exposure (n = 675) versus ADT exposure (n = 458). (I) No ARPI exposure (n = 796) versus ARPI exposure (n = 20). **(J)** Castration-sensitive (n = 675) versus castration-resistant disease (n = 483). **(K)** Normal prostate (n = 173), primary tumor (n = 708), and mCRPC (n = 484). **(L)** Normal prostate (n = 173), primary tumor (n = 708), mARPC (n = 428), mDNPC (n = 22), and mNEPC (n = 34). **(M)** Alive (n = 593) versus deceased (n = 106) at last follow-up. Statistical significance is indicated by asterisks: \*\**P* < 0.01, *** *P* < 0.001, **** *P* < 0.0001; ns, not significant. Statistical tests and multiple-testing correction are indicated in the Methods.

### AXL is Progressively Downregulated During Prostate Cancer Progression, in Contrast to Canonical Oncogenic Drivers MYC and EZH2

We next examined *AXL* expression across PC progression (Fig. 1B). *AXL* mRNA was highest in normal prostate tissue (n = 52), significantly reduced in primary tumors (n= 539; *P* < 0.001), and further reduced in metastatic lesions (n = 100; *P* < 0.001). *AXL* loss therefore tracks monotonically with advanced disease stage. This trajectory was the inverse of that observed for canonical drivers of PC progression profiled in the same cohorts. Consistent with disease progression, the cellular proliferation marker gene *MKI-67* (Supplemental Fig. 1B) and oncogene *MYC* were significantly upregulated from normal prostate to primary tumors and increased further in metastatic disease (Fig. 1C). Likewise, expression of the epigenetic regulator *EZH2*, a key driver of aggressive and neuroendocrine PCa, increased progressively from normal prostate to primary tumors and reached its highest levels in metastatic disease (Fig. 1D). The reciprocal expression patterns of *AXL* and the established PC progression markers *MYC* and *EZH2* across the same patient cohorts argue against a global normalization or batch effect. Instead, these findings support *AXL* downregulation as a biological feature associated with PC progression rather than a technical artifact.

### AXL loss associates with treatment exposure and castration resistance

To define the relationship between *AXL* and PC progression, we first analyzed transcriptomes spanning normal prostate, primary tumor, and metastatic disease, including mARPC, mDNPC, and mNEPC phenotypes. Principal component analysis organized these disease states along a continuum, with *AXL* expression progressively declining along this trajectory and reaching its lowest levels in therapy resistant, metastatic phenotypes (Supplemental Fig. 1C, D). These findings support a progressive loss of *AXL* across the transcriptional landscape of PC. We next stratified *AXL* expression across clinically annotated subsets within an expanded PC cohort from the Prostate Cancer Atlas (Fig. 1E-M). Patients with any prior treatment exposure (n = 489) had significantly lower *AXL* expression than treatment-naive patients (n = 92; *P* < 0.0001, Fig. 1E). Consistent with this, prior taxane exposure (n = 21 versus n = 894; *P* < 0.01, Fig. 1F), androgen deprivation therapy (ADT) exposure (n = 458 versus n = 675; *P* < 0.0001, Fig. 1H), and androgen receptor pathway inhibitor (ARPI) exposure (n = 20 versus n = 796; *P* < 0.001, Fig. 1I) were each associated with reduced *AXL* expression. Notably, radiation therapy (RT) was the only clinical variable associated with increased *AXL* expression. Irradiated patients (n = 552) showed significantly higher *AXL* expression than non-irradiated patients (n = 99; *P* < 0.0001, Fig. 1G), suggesting an RT-associated induction of *AXL* distinct from the effects of systemic ADT or chemotherapy. Consistent with disease progression, *AXL* expression was significantly lower in castration-resistant tumors (n = 483) than in castration-sensitive tumors (n = 675; *P* < 0.0001, Fig. 1J). Across disease states, *AXL* expression also declined from normal prostate tissue (n = 173) to primary tumors (n = 708; *P* < 0.0001) and mCRPC (n = 484; *P* < 0.0001, Fig. 1K), independently corroborating the progressive *AXL* loss observed in Fig. 1B. Stratification of mCRPC by molecular subtype further revealed progressive *AXL* loss with aggressive disease phenotypes and lineage plasticity, as shown in Fig. 1L. *AXL* expression was reduced in mARPC (n = 428; *P* < 0.0001), with the lowest levels in mDNPC (n = 22; *P* < 0.0001) and mNEPC (n = 34; *P* < 0.0001). Thus, *AXL* loss is most pronounced in the increasingly AR low expressing and lineage-plastic states of advanced PC. Patients who died during the study period (n = 106) had significantly lower *AXL* expression than those who remained alive (n = 593; *P* < 0.0001, Fig. 1M), supporting an association between low *AXL* expression and adverse clinical outcome. Collectively, these analyses establish progressive *AXL* loss during PC progression, with the lowest expression in aggressive metastatic disease, and support an association between *AXL* loss and mortality.

### Low AXL Expression is Associated with Reduced Overall Survival in Prostate Cancer

To determine whether the loss of *AXL* observed during PCa progression (Fig. 1) has prognostic significance, we performed Kaplan-Meier survival analysis of the Prostate Cancer Atlas cohort (n = 661 patients with survival annotation), stratifying patients into high- and low-*AXL* groups using the median expression value (log2 CPM = 4.73) as the cutoff (Fig. 2A). This produced two comparably sized groups: *AXL*-low (n = 318) and *AXL*-high (n = 343). The *AXL*-low group had substantially more deaths than the *AXL*-high group; 80 versus 26 deaths. Survival curves separated early, within the first 25 months, and remained distinct through approximately 110 months, with the *AXL*-low group showing consistently lower survival probability across this interval. At 100 months, estimated survival was approximately 0.62 in the *AXL*-low group compared with 0.70 in the *AXL*-high group. Median overall survival was 115.1 months in the *AXL*-low group and 132.8 months in the *AXL*-high group. Low *AXL* expression was associated with significantly worse overall survival (HR = 0.51, 95% CI 0.32-0.80, p = 0.003 for high versus low expression), indicating that patients with low *AXL* expression had a higher risk of death during follow-up than patients with high *AXL* expression. The confidence interval remained below unity, supporting a significant association between low *AXL* expression and adverse outcome. Beyond approximately 110 months, the survival curves converge and cross near 145-150 months, coinciding with markedly reduced numbers of patients at risk and widening confidence intervals. At these late follow-up intervals, only 13 *AXL*-low and 3 *AXL*-high patients remained at risk at 150 months, decreasing to 6 and 2, respectively, at 200 months. Therefore, survival estimates at these time points should be interpreted cautiously. The prognostic association was primarily driven by the first 110 months of follow-up, when the majority of events and patients at risk were observed.

**Figure 2.**
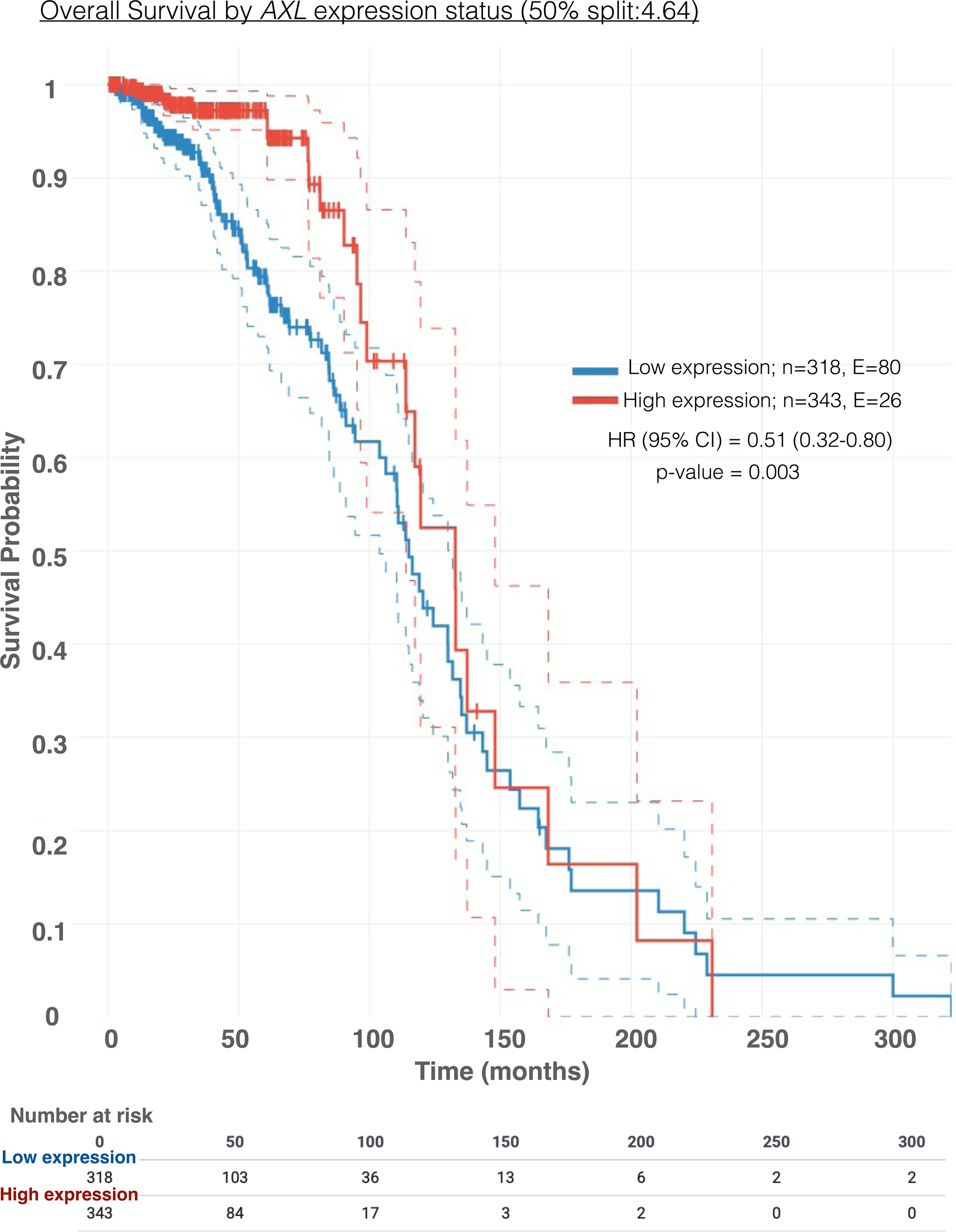
Low AXL expression is associated with reduced overall survival in prostate cancer. (Low AXL expression is associated with reduced overall survival in prostate cancer. **(A)** Kaplan-Meier estimates of overall survival in the Prostate Cancer Atlas cohort (n = 661 patients with survival annotation), stratified by median *AXL* transcript expression (ENST00000301178.9; log_2_ CPM = 4.73). Patients were classified as *AXL*-low (blue; n = 318, 80 deaths) or *AXL*-high (red; n = 343, 26 deaths). Solid lines represent Kaplan-Meier survival estimates, dashed lines indicate 95% confidence intervals, and tick marks denote censored observations. Median overall survival was 115.1 months in the *AXL*-low group and 132.8 months in the AXL-high group. The hazard ratio for high versus low *AXL* expression was 0.51 (95% CI, 0.32-0.80; *P* = 0.003; Wald test). Numbers at risk are shown at 50-month intervals. The risk table beneath the plot reports the number of patients remaining at risk in each group at 50-month intervals. Note the marked attrition after 150 months (13 *AXL*-low and 3 *AXL*-high at risk), which accounts for the widening confidence bands and curve convergence at late time points.

Because the curves cross at late time points, the hazard ratio represents an average effect concentrated in early follow-up; HR (95% CI) = 0.51 using Cox proportional hazards regression model. These findings are consistent with the clinical stratification in Fig. 1M, where deceased patients exhibited significantly lower *AXL* expression than surviving patients, together supporting an association between *AXL* loss and adverse patient outcome.

Next, we asked whether genomic alterations in *AXL* showed a similar clinical association. *AXL* alterations were rare but clinically consequential. In the TCGA-PRAD cohort, 21 of 2,201 patients (∼1%) harbored an *AXL* alteration, i.e., loss-of-function mutations or copy loss, and these patients had significantly shorter overall survival than those with unaltered *AXL* (log-rank *P* = 0.0394; Supplemental Fig. 1E). Survival curves separated early and continued to diverge, with approximately 45% of patients with *AXL* alterations surviving at 55 months compared with ∼60% of those without alterations; no patient with an *AXL* alteration survived beyond 82 months. Given the small number of cases with *AXL* alterations, these findings should be considered hypothesis generating rather than definitive. Nevertheless, the concordant association of low *AXL* expression and *AXL* alterations with shorter survival further supports the clinical relevance of *AXL* loss in PCa. Together, these clinical datasets establish *AXL* loss as a feature associated with poor patient outcome and provide the rationale for functional interrogation of *AXL* loss in PCa models.

### AXL Loss Enhances Proliferation and Migration

The clinical data in Fig. 1 and 2 indicate that *AXL* expression progressively decreases with PCa progression and is associated with adverse clinical outcomes, with the lowest levels observed in mDNPC and mNEPC subtypes. To investigate the functional consequences of *AXL* loss, we first identified a model with sufficient endogenous *AXL* expression to enable isogenic deletion, as genetic loss-of-function studies are not informative in models with little or no baseline receptor expression. We therefore examined *AXL* and *GAS6* expression across a panel of PCa cell lines (Supplemental Fig. 2A). *AXL* was predominantly expressed in cell lines that are of basal/intermediate lineage, with the highest levels in DU145, PC3, and PC3M, whereas AR expressing cell lines and the NEPC line NCI-H660 expressed little or no *AXL*. The ligand *GAS6* was broadly expressed across the panel, indicating that *AXL* expression primarily determined the presence of the GAS6-AXL signaling axis. We selected PC3, a bone metastasis derived cell line, for subsequent studies. PC3 cells expressed both AXL and GAS6, establishing an endogenous receptor-ligand axis suitable for genetic disruption. PC3 has been classified as mDNPC in some studies and reported to exhibit mNEPC features in others [44, 45], consistent with its use as a model of aggressive metastatic PCa. To test whether AXL loss has functional consequences beyond its correlation with advanced PCa, we generated AXL-deficient PC3 cells by CRISPR-Cas9 editing and assessed proliferation and migration. AXL knockout efficiency was assessed by immunoblotting (Supplemental Fig. 2B) and flow cytometric quantification of cell-surface AXL protein using a PE-conjugated anti-AXL antibody (Fig. 3A). We established the background threshold using parental PC3 cells expressing a non-targeting sgRNA (*AXL*^WT^; sgROSA) stained with an isotype-matched IgG control, which showed no detectable PE signal (Fig. 3A, left panel). In contrast, cells stained with the anti-AXL antibody showed a clear AXL-positive population, confirming abundant cell surface AXL expression in PC3 cells (*AXL*^WT^); Fig. 3A, middle panel. Whereas AXL knockout cells (PC3 sgAXL; *AXL*^KO^) stained with the same antibody remained within the AXL-negative population, demonstrating near-complete loss of AXL protein; see Fig. 3A, right panel.

**Figure 3.**
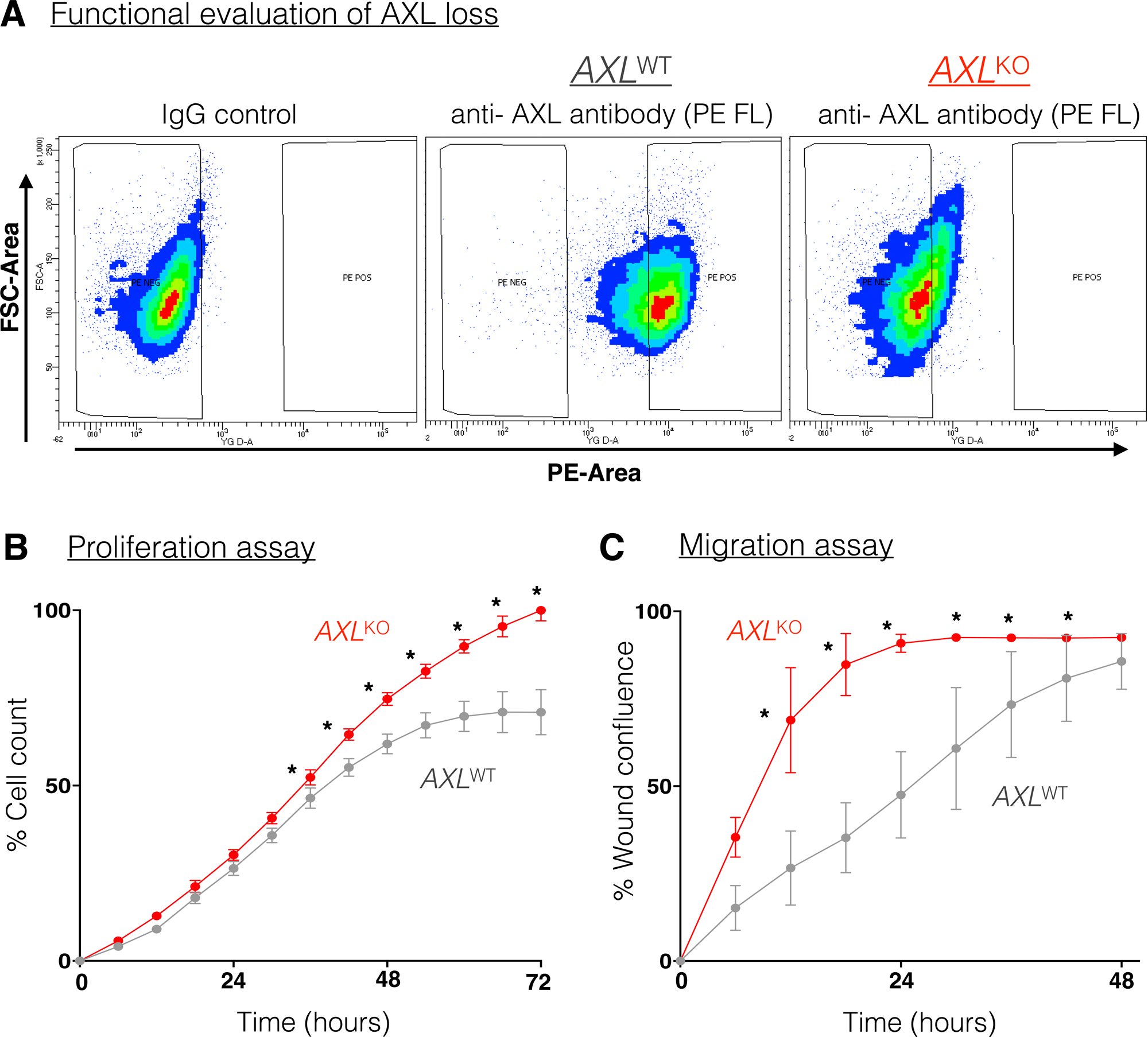
AXL loss increases proliferation and migration in PC3 prostate cancer cells. **(A)** Flow cytometric validation of AXL knockout. Density plots show forward scatter area (FSC-Area) versus PE fluorescence area (PE-Area) for PC3 cells. Left: PC3 *AXL*^WT^ cells stained with isotype-matched IgG control, defining the PE-negative background. Center: PC3 *AXL*^WT^ cells stained with PE-conjugated anti-AXL antibody, showing a distinct PE-positive population. Right: PC3 *AXL*^KO^ cells stained with the same anti-AXL antibody, showing loss of the PE-positive population. Gates denote PE-negative (PE NEG) and PE-positive (PE POS) regions; color scale reflects event density. **(B)** Proliferation of *AXL*^WT^ (grey) and *AXL*^KO^ (red) PC3 cells over 72 hours, expressed as percent cell count normalized to the maximum value. **(C)** Wound-closure migration assay for *AXL*^WT^ (grey) and *AXL*^KO^ (red) PC3 cells over 48 hours, expressed as percent wound confluence. Points represent the mean ±SD. Statistical significance is indicated by asterisks: \**P* < 0.05, \*\**P* < 0.01, *** *P* < 0.001, **** *P* < 0.0001; ns, not significant. Statistical test is two-way repeated-measures ANOVA with Sidak’s multiple-comparisons test

Next, we examined the impact of AXL loss on PC3 cell proliferation. Longitudinal cell counts over 72 hours revealed comparable growth between *AXL*^WT^ and *AXL*^KO^ cells during the first 24 hours (Fig. 3B). Beginning at 24 hours, *AXL*^KO^ cells showed significantly greater cell growth compared to *AXL*^WT^ cells, with differences maintained at subsequent time points (*P* < 0.05). At the 72 hour timepoint, *AXL*^KO^ cells exhibited higher normalized cell counts than *AXL*^WT^ cells. Together, our data demonstrate that loss of *AXL* alters PC3 growth kinetics, resulting in increased cell growth over time. Our finding that AXL loss enhances PC3 cell proliferation appears to conflict with Axelrod et al., who reported no proliferative change in AXL-null PC3luc cells generated using the same exon 2 gRNA [46]. This discrepancy may reflect differences in assay duration; whereas the previous study measured BrdU incorporation over 24 hours, our longitudinal cell counts showed no difference between *AXL*^WT^ and *AXL*^KO^ cells during the first 24 hours, with divergence emerging only thereafter (Fig. 3B). Thus, the earlier assay may have been insufficiently long to detect the delayed proliferative phenotype. Differences in clonal architecture may also contribute, as the previous study examined two single-cell derived knockout clones and one control clone [46], potentially introducing clone-specific effects. Notably, the two knockout clones responded discordantly to GAS6 stimulation, underscoring the potential influence of clonal variability [46].

Given that metastatic dissemination requires both sustained tumor cell expansion and acquisition of migratory capacity [47–49], we next examined whether *AXL* loss alters the migratory phenotype of PC3 cells. Migratory capacity was assessed by a wound-closure assay over 48 hours (Fig. 3C). The migratory difference between genotypes emerged early. At 6 hours, *AXL*^KO^ cells had closed approximately 35% of the wound compared with approximately 15% for *AXL*^WT^ cells. By 12 hours, *AXL*^KO^ cells had closed approximately 70% of the scratch wound compared with approximately 25% for *AXL*^WT^ cells (*P* < 0.05). *AXL*^KO^ cells achieved near complete closure (>90%) by 24 hours, whereas *AXL*^WT^ cells reached comparable wound closure only near the end of the 48 hour assay. Significant differences between genotypes were observed at 6, 12, 18, 24, 30, 36, and 42 hours, with the curves converging by 48 hours as both populations approached complete wound closure. Together, these findings demonstrate that *AXL* loss enhances both proliferative capacity and migratory behavior in PC3 cells, supporting a functional role for *AXL* loss in promoting aggressive cellular phenotypes.

### AXL Loss Creates a Therapeutically Exploitable Kinase Dependency

Having established that *AXL* loss enhances proliferation and migration in PC3 cells (Fig. 3), we asked whether *AXL*^KO^ cells acquire pharmacologically exploitable vulnerabilities. Genetic or pharmacologic loss of a receptor tyrosine kinase rarely occurs in isolation; cells typically compensate by rewiring parallel survival pathways, and this adaptation frequently generates collateral dependencies to pathways that were dispensable in the parental state [29, 35]. We therefore performed an unbiased high-throughput screen of a 1,814 compound annotated kinase inhibitor library (Selleckchem) in isogenic *AXL*^KO^ and *AXL*^WT^ cells to identify pharmacological vulnerabilities associated with AXL loss; schematic in Supplemental Fig. 2C. Each compound was evaluated at 0.5 μM and 2.5 μM in three biological replicates, and cell viability was quantified by MTT assay following 72 hours of treatment across 22 assay plates. The optical density (OD) value measured from each well was adjusted relative to the DMSO or water vehicle control wells on the same plate to account for plate-to-plate and assay variation. Compound activity was quantified using strictly standardized mean difference (SSMD), a statistical metric widely used for high-throughput screening to identify compounds producing robust effects relative to assay variability [50, 51] schematic in Supplemental Fig. 2D. SSMD values were calculated for each compound, genotype, and concentration relative to the corresponding vehicle control distribution. Of the compounds screened, 114 met the predefined responder threshold (SSMD <-3) in *AXL*^KO^ cells at one or both concentrations (Supplemental Fig. 3). To determine whether these hits converged on specific signaling programs, we extracted pathway annotations from the drug metadata provided by SelleckChem and the pathway composition of the responder set was compared with that of the full library (Fig. 4 and Table1). Hits were distributed across 22 annotated pathways but were not uniformly represented. Compounds targeting the cell cycle regulators preferentially reduced the viability of *AXL*^KO^ cells (data not shown). PI3K/AKT/mTOR inhibitors constituted the largest class of *AXL*^KO^ selective hits (n=12 of 27 compounds; 44%), exceeding the next most represented pathways, protein tyrosine kinase and TGF-β/Smad signaling. Relative to their representation in the library (n= 213 compounds, 5.3%) PI3K/AKT/mTOR inhibitors were enriched 4-fold among responders (*P* adj = 0.0004, Fishers exact test with FDR correction, Supplemental Fig. 4A). Several pathways well represented in the library, including GPCR and G protein signaling, epigenetics, and cell cycle, yielded few or no selective hits despite comparable or greater library coverage.

**Figure 4.**
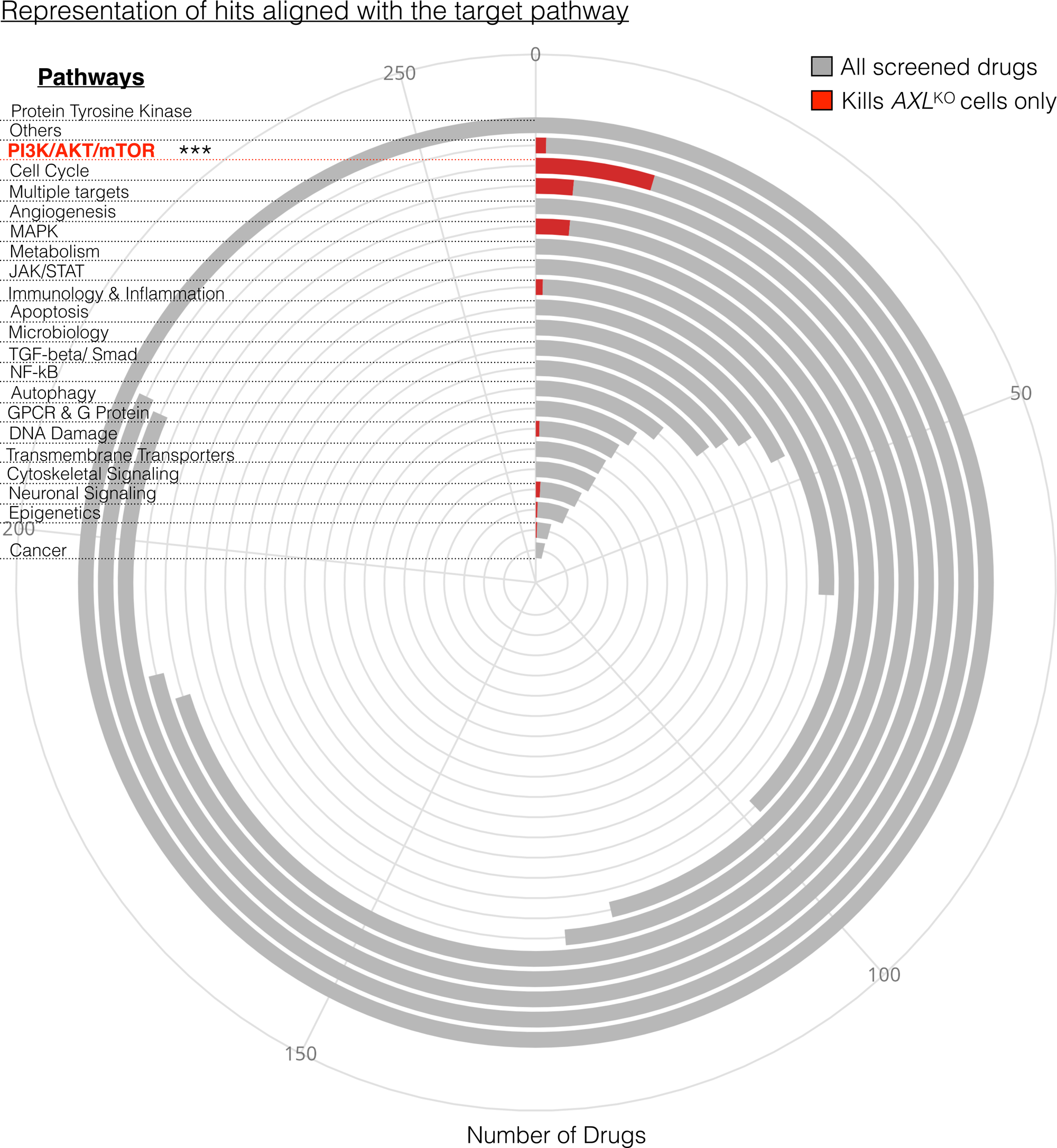
Kinase inhibitor screen reveals preferential sensitivity of *AXL*^KO^ PC3 cells to PI3K/AKT/mTOR pathway inhibition. **(A)** Radial bar plot showing the distribution of screened compounds across annotated target pathways. Grey bars indicate the total number of compounds assigned to each pathway, and red bars indicate the number of compounds within each pathway that induced cell killing in *AXL*^KO^ PC3 cells, as defined by the prespecified hit criteria (SSMD < −3 at both concentrations). The radial axis represents compound counts, with gridlines at 50, 100, 150, 200, and 250. The PI3K/AKT/mTOR pathway, highlighted in red text, yielded the highest number of hits (12 compounds; \*\*\**P* < 0.0001). Pathway-level enrichment of hits relative to library representation was assessed using Fisher’s exact test with FDR correction for multiple comparisons across 22 pathways.

The convergence of *AXL*^KO^ selective compounds on PI3K/AKT/mTOR signaling suggests that loss of *AXL* creates increased reliance on this pathway for cellular survival.

### Kinase Screen Hits validate as selective vulnerabilities of *AXL*^KO^ cells and converge on mTORC1 signaling

The shortlisted compounds that met the hit criterion of SSMD < −3 at both concentrations are shown in Fig. 5A. We performed unsupervised clustering to distinguish DMSO vs drug treated replicates at both doses and found that all identified hits reduced viability relative to vehicle controls. The hits encompassed multiple signaling pathways, with predominantly PI3K/AKT/mTOR pathway inhibitors. These included AZD8186, taselisib, pictilisib, PF-4989216, torin 1, torin 2 and zotarolimus. Additional kinase targets included ATR (VE-822), Aurora A (MK-8745), PDK1 (GSK2334470), PAK4 (PF-3758309), BTK (CGI1746) and TTK/MPS1 (BAY 1217389), alongside the multikinase inhibitors GNF-7, CX-6258, XL228 and dasatinib. The non-kinase-specific compounds deguelin, MCB-613, bardoxolone methyl, and cinobufagin also met the hit criteria. Cinobufagin produced the largest SSMD magnitude at both concentrations (−25.62 at 0.5 µM and −7.25 at 2.5 µM). Notably, hit magnitude did not consistently increase with concentration. Sixteen of the 22 compounds showed greater SSMD magnitudes at 0.5 µM than at 2.5 µM, whereas deguelin, GNF-7, dasatinib, pictilisib and bardoxolone methyl showed stronger effects at the higher concentration. Bardoxolone methyl exhibited the largest increase in SSMD magnitude, from −4.30 to −9.22.

**Figure 5.**
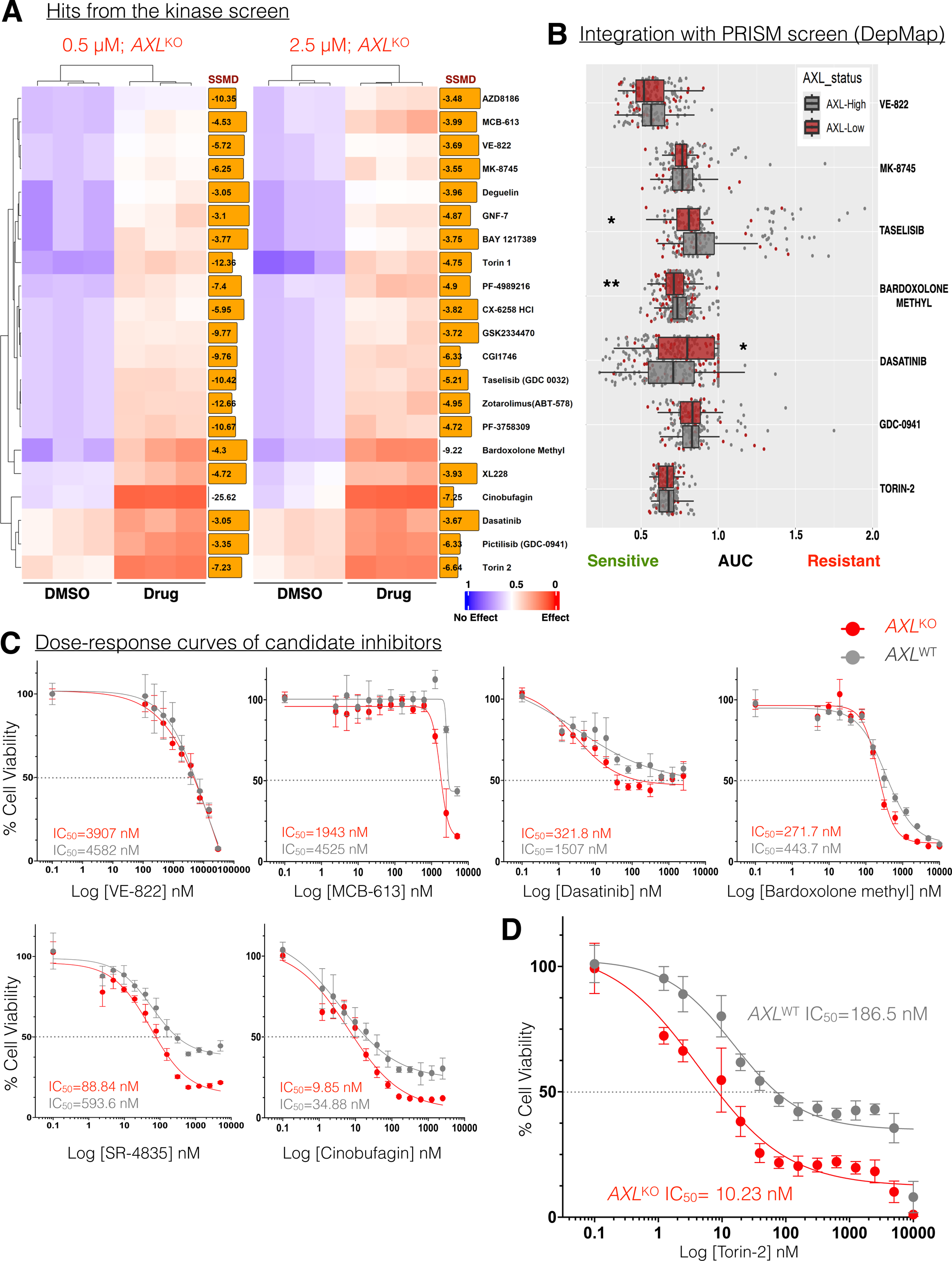
AXL loss sensitizes PC3 cells to inhibition of mTOR signaling. **(A)** Heatmap of the 22 compounds meeting the hit criterion in the kinase inhibitor screen in *AXL*^KO^ PC3 cells at 0.5 µM (left) and 2.5 µM (right). Columns represent DMSOvtreated controls and compound-treated replicates, and rows represent individual compounds. Colors indicate optical density normalized to the mean of the vehicle controls on each replicate plate, ranging from blue (no effect) through white to red (effect). Boxed values to the right of each panel indicate the SSMD for each compound; more negative values indicate greater reductions in viability relative to vehicle controls. Compounds were classified as hits if SSMD < −3 at both concentrations. **(B)** Sensitivity of AXL-low and AXL-high cancer cell lines to seven hit compounds identified in the kinase screen and evaluated in the PRISM drug repurposing screen (DepMap, 20Q2): bardoxolone methyl, taselisib, dasatinib, Torin-2, pictilisib (GDC-0941), VE-822, and MK-8745. *AXL*-low cell lines were significantly more sensitive to taselisib and bardoxolone methyl, whereas *AXL-* high cell lines were more sensitive to dasatinib. Cell lines were stratified by *AXL* expression (DepMap Public 24Q4) into *AXL*-Low (first quartile, log_2_ (TPM + 1) < 0.4; red; n = 40) and *AXL-*High (third quartile, log_2_ (TPM + 1) > 6.4; gray; n = 155) groups. Boxplots show the area under the dose-response curve, with lower AUC values indicating greater drug sensitivity. Each point represents one cell line. For each drug, AUC values were converted to drug-specific z-scores, and cell lines with z-scores < −1 were classified as sensitive. The proportions of sensitive cell lines were compared between groups using two-sided Fisher’s exact tests. \**P* < 0.05, \*\**P* < 0.01; ns, not significant. *P* values were not adjusted for multiple comparisons. **(C)** Dose-response viability curves for six compounds preferentially active following AXL loss (SR-4835, dasatinib, cinobufagin, MCB-613, uprosertib, and bardoxolone methyl) in *AXL*^WT^ (gray) and *AXL*^KO^ (red) PC3 cells following 72 h of treatment. **(D)** Dose-response viability curves for Torin-2 in *AXL*^WT^ (gray) and *AXL*^KO^ (red) PC3 cells. Points represent mean ±SD. The IC_50_ values were 10.23 nM for *AXL*^KO^ cells and 186.5 nM for *AXL*^WT^ cells. In **(C)** and **(D)**, viability is normalized to vehicle-treated controls, and data are presented as mean ± SD from 2-3 biological replicates. Curves were fitted by four parameter logistic nonlinear regression (GraphPad PRISM) and the half maximal values (IC_50_) with 95% confidence intervals were interpolated from the fitted curves.

To determine whether these vulnerabilities were recapitulated in an independent dataset, we interrogated the PRISM repurposing screen in DepMap (n = 1,448 compounds; Fig. 5B) and stratified cell lines by *AXL* expression. An unbiased analysis across all 1,448 PRISM compounds was performed to identify compounds with preferential activity in *AXL*-low versus *AXL*-high expression cell lines. In Fig. 5B, we highlight seven hits from the kinase screen that were also identified in the unbiased PRISM database analysis. The *AXL*-low cell lines (n = 40 cell lines) were significantly more sensitive than *AXL*-high cell lines (n = 155 cell lines) to bardoxolone methyl (*P* < 0.01) and taselisib (*P* < 0.05). In contrast, our analysis showed that *AXL*-low cell lines in the PRISM dataset are resistant to dasatinib (*P* < 0.05). Torin-2, GDC-0941, VE-822, and MK-8745 showed the same directional trend but did not reach statistical significance. Because PRISM profiles a cross-lineage panel rather than isogenic pairs, these comparisons reflect correlation with endogenous *AXL* expression rather than the consequence of *AXL* deletion. Nonetheless, their concordance with the isogenic screen supports an association between low *AXL* expression and sensitivity to these agents. These findings identify compounds that reduce viability in *AXL*^KO^ PC3 cells but do not establish whether their activity is selective for AXL loss. To assess genotype selectivity, we performed preliminary dose-response validation of twelve candidate compounds selected from the Kinase screen and the PRISM analysis in isogenic *AXL*^KO^ and *AXL*^WT^ cells (Supplemental Fig. 5A). These were selected from the 114 compounds that met the responder threshold (SSMD< −3) in *AXL*^KO^ cells, prioritized by *AXL*^KO^ selectivity in the screen, concordance with the PRISM analysis and pathway representation. Compound potency varied by more than two orders of magnitude: cinobufagin and Torin-2 had single-to low double-digit nanomolar IC_50_ values in *AXL*^KO^ cells, whereas the nilotinib IC_50_ exceeded 5,000 nM in both genotypes, precluding estimation of a fold-shift (Supplemental Fig. 5A). To quantify selectivity independently of absolute potency, we calculated the IC_50_ fold shift (*AXL*^WT^ / *AXL*^KO^) for 11 compounds and classified compounds with a fold shift > 1.3 and Benjamini-Hochberg adjusted p<0.05 as selective for AXL loss (Supplemental Fig. 5C). Eight compounds met both selectivity criteria: Torin-2 (18.2-fold; IC_50_, 10.23 vs. 186.5 nM in *AXL*^KO^ and *AXL*^WT^ cells, respectively), SR-4835 (6.68-fold; 88.84 vs. 593.6 nM), dasatinib (4.68-fold; 321.8 vs. 1,507 nM), cinobufagin (3.54-fold; 9.85 vs. 34.88 nM), MCB-613 (2.33-fold; 1,943 vs. 4,525 nM), uprosertib (2.29-fold; 359.9 vs. 825.1 nM), bardoxolone methyl (1.63-fold; 271.7 vs. 443.7 nM), and SGC-GAK-1 (1.30-fold; 898.8 vs. 1,171 nM). In contrast, taselisib (1.02-fold), tomatine (1.16-fold), and VE-822 (1.17-fold) showed minimal differences in potency between genotypes. Because the SGC-GAK-1 dose-response curves intersected within the tested concentration range, complicating interpretation of its IC_50_ fold-shift, the remaining seven selective compounds were selected for further analysis, with their dose-response curves shown in Fig. 5C and D.

Torin-2 exhibited the greatest genotype selectivity (Fig. 5D). In addition to its 18.2-fold IC_50_ shift, observed viability in *AXL*^KO^ cells remained below 25% at concentrations above 100 nM, compared with approximately 40% in *AXL*^WT^ cells. Similarly, SR-4835 and cinobufagin produced lower observed viability at the highest concentrations tested in *AXL*^KO^ cells than in *AXL*^WT^ cells (Fig. 5C). These findings suggest that AXL loss influences not only drug potency but also the extent of growth inhibition achieved at the concentrations tested. Importantly, genotype selectivity did not simply reflect absolute potency: cinobufagin was the most potent compound in *AXL*^KO^ cells but ranked fourth in selectivity, whereas Torin-2 was less potent yet exhibited the greatest genotype selectivity.

Within the PI3K/AKT/mTOR signaling pathway, both the mTOR inhibitor Torin-2 and the AKT inhibitor uprosertib showed selectivity for *AXL*^KO^ cells, whereas the PI3Kα selective inhibitor taselisib did not, despite being identified as a hit in both the isogenic screen and the PRISM analysis. Together, these findings suggest that AXL loss may preferentially sensitize PCa cells to inhibition of the AKT/mTOR axis and identify mTOR inhibition, represented by Torin-2, as a prominent pharmacological vulnerability associated with AXL loss.

## Discussion

The published literature describes a context dependent role of AXL in PCa, implicating it both as an oncogenic driver of tumor progression and a tumor suppressor that promotes DTC dormancy [9, 12, 13, 52–57]. Additional studies have linked *AXL* to other aspects of tumor biology, including treatment response and interactions with the tumor microenvironment [9, 12, 13, 57]. Thus, the consequences of *AXL* expression may depend on the biological context in which it functions. This context-dependent biology provides a framework for our findings. We profiled *AXL* expression across PCa disease stages and found that its loss is associated with aggressive disease. Functionally, *AXL* deletion enhanced proliferation and migration, prompting us to hypothesize that *AXL* deficiency creates compensatory signaling dependencies that may be therapeutically exploitable. To test this hypothesis, we performed an unbiased kinase inhibitor screen in isogenic *AXL*^WT^ and *AXL* ^KO^ cells and identified signaling dependencies associated with the AXL-deficient state. To our knowledge, this is the first study to systematically characterize drug-response dependencies associated with AXL loss in PCa. Together, our findings link AXL loss to aggressive disease and identify potential therapeutic vulnerabilities in AXL-deficient cells.

Our analysis of clinical datasets supports an association between *AXL* loss and PCa progression. *AXL* expression declined from normal prostate to primary and metastatic PCa and was lowest in metastatic double-negative and neuroendocrine disease. Conversely, within the same cohort, *MYC*, *EZH2*, and the cell proliferation marker *MKI67* increased with progression, consistent with more proliferative and lineage-plastic tumor phenotypes. Importantly, low *AXL* expression was also associated with shorter overall survival, whereas high expression was associated with a longer overall survival. Our findings extend the clinical observations across additional disease states and treatment contexts.

Radiotherapy was an exception to the overall association between treatment exposure and lower *AXL* expression, with irradiated patients showing higher expression than non-irradiated patients (n = 552 versus n = 99; *P* < 0.0001). This finding is consistent with reports linking *AXL* to DNA repair and radioresistance [58–60], although our analysis cannot determine whether radiotherapy directly induces *AXL* or whether this association reflects differences in treatment history or disease characteristics. More broadly, hypoxia and differences in the cellular composition of metastatic tissues may influence measured *AXL* expression and activity [53, 57]. These considerations highlight the limitations of bulk transcriptomic analyses, which cannot establish tumor-cell-specific changes in *AXL* signaling. Finally, our analysis of the clinical data establishes associations rather than causality and does not establish *AXL* as an independent prognostic factor without multivariable analyses accounting for relevant clinical and molecular covariates. The low frequency of *AXL* alterations in the analyzed genomic dataset also limits interpretation of mutation-based survival comparisons.

Beyond its association with disease progression, our findings suggest that exploiting therapeutic vulnerabilities associated with *AXL* loss may provide a strategy to target proliferative cells emerging from otherwise undetectable and difficult to target dormant DTCs. In the kinase inhibitor screen, Torin-2 showed an approximately 18.2-fold lower IC_50_ in *AXL*^KO^ than *AXL*^WT^ cells. Similarly, Uprosertib, an AKT inhibitor, showed modest selectivity (2.29-fold), whereas the PI3Kα-selective inhibitor taselisib showed essentially no genotype selectivity. The differential responses to inhibitors acting at distinct levels of the pathway provide an additional clue to the nature of this vulnerability. These differences suggest that the observed vulnerability is not a uniform consequence of inhibiting the PI3K/AKT/mTOR pathway but may instead reflect a more specific dependency at or downstream of AKT/mTOR signaling in the absence of AXL. Further mechanistic studies are needed to determine whether *AXL* loss directly alters mTORC1 activity and whether this pathway mediates the observed drug sensitivity.

Although the DepMap PRISM analysis did not independently validate a significant association between *AXL* expression and Torin-2 response, it provides a complementary resource for exploring drug-response relationships associated with *AXL* expression. Our findings suggest that AXL-low cells may have distinct signaling dependencies rather than being broadly sensitive to drug treatment. However, cross cell-line comparisons may be confounded by genomic background, tissue lineage, and other molecular features. The DepMap PRISM analysis identified an association between *AXL* expression and taselisib response. However, taselisib showed no genotype selectivity in our isogenic PC3 models, suggesting that this association may reflect differences between cell lines rather than AXL status itself. Further work is needed to determine whether *AXL* loss itself drives these responses or serves as a marker of a broader cellular state.

Our study also highlights that a primary-screen hit represents an initial identification of potential therapeutic targets rather than a validated therapeutic vulnerability, and that multiple orthogonal approaches are required to establish genotype-selective sensitivity. Of the 114 compounds that reduced viability in *AXL*^KO^ cells, only a small subset showed concordant activity in the independent PRISM dataset, and subsequent dose-response analysis revealed that activity in both datasets did not necessarily translate into AXL dependent selectivity, as exemplified by taselisib. These discrepancies are consistent with known limitations of drug-screening approaches, including assay interference and compound-specific artifacts in single-concentration screens [61], substantial variability in drug-sensitivity measurements across pharmacogenomic datasets [62] and confounding of endpoint-based metrics such as IC_50_ by differences in cell proliferation rates [63]. The latter limitation is particularly relevant because *AXL*^KO^ cells proliferate faster than *AXL*^WT^ cells. Accordingly, integrating an isogenic primary screen, cross-cell-line PRISM analysis, and quantitative dose-response fold-shift analysis provided a more robust framework for prioritizing Torin-2 than any single assay alone.

Inhibitors with reported activity against TAM-family receptors did not emerge as hits in *AXL*^KO^ cells, providing no evidence of an acute compensatory dependency on residual TAM signaling. However, this finding is inconclusive given the limited concentration range, unconfirmed target engagement, and unknown baseline MERTK and TYRO3 expression in PC3 cells. Ligand-dependent or microenvironmentally driven compensation and longer-term adaptation may not be captured by this screen. Defining the roles of MERTK and TYRO3 in adaptation to AXL loss will require genetic approaches, including single and combinatorial knockout in the *AXL*^KO^ background, coupled with assessment of GAS6 stimulated signaling and dormancy associated phenotypes in vivo.

We provide a roadmap for investigating how *AXL* loss shapes therapeutic dependencies across different molecular contexts of PCa. PC3 cells are *PTEN*-null, *TP53*-null, and AR-independent [44, 45], providing a relevant model of aggressive disease. Because *PTEN* loss alters PI3K/AKT signaling, extending these findings to additional isogenic models with different *PTEN* and *TP53* backgrounds and to patient-derived PCa models will help define the contexts in which *AXL* loss confers this vulnerability. The increased sensitivity to Torin-2 provides a rationale for evaluating clinically investigated mTOR inhibitors, including sapanisertib and vistusertib, in *AXL*-deficient PCa models. These studies will help establish the molecular contexts in which *AXL* loss-associated vulnerabilities can be exploited therapeutically.

In summary, our study identifies *AXL* loss as a feature of PCa progression and demonstrates that *AXL* low status is associated with aggressive disease and poor overall survival. We further identify a potential therapeutic vulnerability of *AXL*^KO^ cells to PI3K/AKT/mTOR pathway inhibition. Together, our findings link *AXL* loss to aggressive disease and altered drug sensitivity, providing new insight into the context-dependent roles of AXL in PCa progression.

## Data Availability

Cell lines and models described in this study are available upon request through the Mathew Laboratory, subject to applicable UNMC material transfer agreements and institutional approvals.

## Code Availability

The software and algorithms for data analyses used in this study are published and referenced throughout the methods section. This study does not report original code. Any additional information required to reanalyze the data reported in this paper is available from the lead contact upon request.

## Authors Contributions

Conceptualization, S.S., K.M., S.K., A.N., D.K., and G.M.; methodology, S.S., K.M., S.K., A.N., D.K., and G.M.; investigations, S.S., K.M., and G.M.; formal analyses, S.S., K.M., and G.M. data curation, S.S., K.M., and G.M.; resources, G.M.; writing, S.S., K.M., and G.M.; funding acquisition, G.M.

## Funding

This work was supported by the National Institutes of Health/National Cancer Institute, NIGMS COBRE (P20GM121316), R01CA274605, American Cancer Society Institutional Research Grant, and FPBCC pilot grants to G. Mathew, and the APC was funded by the same awards. The Natarajan laboratory was supported by National Cancer Institute R01CA276846.

The authors used Grammarly to improve the quality of the writing for specific sections of the manuscript and BioRender.com to produce the graphical abstract and schematics included in specific figures.

## Acknowledgment

The authors thank all members of the Natarajan, Kelly, and Mathew laboratories for helpful discussions and critical reading of the manuscript.

## Abbreviations

ADT, androgen deprivation therapy; ARPI, androgen receptor pathway inhibitor; CPM, counts per million; IQR, interquartile range; mARPC, metastatic AR-active prostate cancer; mCRPC, metastatic castration-resistant prostate cancer; mDNPC, metastatic double-negative prostate cancer; mNEPC, metastatic neuroendocrine prostate cancer; PRAD, prostate adenocarcinoma; VST, variance-stabilizing transformation.

## FIGURE LEGEND

**Supplementary Figure 1: *AXL* expression across cancer types and prostate cancer progression and its association with patient survival. (A)** *AXL* mRNA expression in normal tissue (green) and primary tumors across 16 TCGA cancer types, shown as log_2_ CPM. Prostate adenocarcinoma (PRAD) is highlighted in bold. Boxes indicate the median and interquartile range (IQR), and whiskers extend to 1.5 x IQR. Sample numbers are shown below each pair of distributions (normal, green; tumor, blue). Statistical comparisons were performed using two-sided Wilcoxon signed-rank test, BH FDR; \*\*\**P* < 0.001; ns, not significant. KIRC, kidney renal clear cell carcinoma; KIRP, kidney renal papillary cell carcinoma; BRCA, breast invasive carcinoma; LUAD, lung adenocarcinoma; STAD, stomach adenocarcinoma; THCA, thyroid carcinoma; HNSC, head and neck squamous cell carcinoma; LUSC, lung squamous cell carcinoma; ESCA, esophageal carcinoma; PRAD, prostate adenocarcinoma; BLCA, bladder urothelial carcinoma; COAD, colon adenocarcinoma; READ, rectum adenocarcinoma; UCEC, uterine corpus endometrial carcinoma; KICH, kidney chromophobe; LIHC, liver hepatocellular carcinoma. **(B)** *MKI67* mRNA expression in normal prostate (n = 52), primary prostate tumors (n = 539), and metastatic prostate cancer (n = 100) from GDC RNA-seq data. Red horizontal lines indicate the median expression; \*\*\*\**P* < 0.0001; two-sided Wilcoxon rank-sum with BH correction. **(C)** Principal component analysis of transcriptomes from the Prostate Cancer Atlas [41], colored by disease stage. Samples include normal prostate, primary tumor, metastatic androgen receptor positive prostate cancer (mARPC), metastatic double negative prostate cancer (mDNPC), and metastatic neuroendocrine prostate cancer (mNEPC). The black line indicates the progression trajectory across disease stages. **(D)** Same PCA projection as in **(C)**, colored according to *AXL* expression normalized log_2_ CPM, with lower expression shown in blue and higher expression in red. **(E)** Kaplan-Meier estimates of overall survival in patients with PCa with (red, n = 21) or without (blue, n = 2,180) *AXL* genomic alterations from the PRAD cohort (all) available through cBioPortal. Tick marks indicate censored patients, and the table below shows the number of patients at risk at each time point. Survival distributions were compared using the log-rank test (p = 0.0394). *AXL* alterations included loss-of-function mutations and copy loss.

**Supplementary Figure 2: *AXL* expression in prostate cancer cell lines, generation of *AXL*-knockout PC3 cells, and design of the kinase inhibitor screen. (A)** Heatmap of *AXL* and *GAS6* mRNA expression in nine PCa cell lines, shown as log2(TPM + 1) from [42]. Values are printed in each cell and shaded from 0 (white) to 10 (black). **(B)** Immunoblot of AXL protein in PC3-Cas9 cells transduced with a control guide targeting the *ROSA26* locus (sg *ROSA*) or with one of three independent guides targeting *AXL* (sg*AXL1-*3). PNT2-C2 cells serve as an AXL-positive control. β-Actin is the loading control. **(C)** Outline of the kinase inhibitor screen. PC3 *AXL*^WT^ and *AXL*^KO^ cells were seeded in 96-well plates on day 0. On day 1, compounds from the SelleckChem kinase inhibitor library (1814 compounds) were added at 0.5 µM or 2.5 µM. Viability was measured by MTT assay on day 4, after 72 h of exposure. Each plate held test compounds in columns 1-11. Column 12 held vehicle (DMSO or water) negative-control wells in rows A-B and E-H, and bemcentinib (10 µM) positive-control wells in rows C-D. **(D)** Hit identification workflow. Each concentration was screened in three replicate plates. Drug-well optical density (OD) was normalized to the mean vehicle OD on the same replicate plate. For each compound, the strictly standardized mean difference was then calculated as SSMD = [mean(drug) − mean(DMSO)] / √[SD(drug)^2^ + SD(DMSO)^2^]. Compounds with SSMD < −3.0 were classified as hits.

**Supplementary Figure 3:** Strictly standardized mean difference (SSMD) values for compounds from the SelleckChem kinase inhibitor library (1814 compounds) screened against *AXL*^WT^ and *AXL*^KO^ cells at 0.5 µM (left) and 2.5 µM (right). Cells were exposed to each compound for 72 h, and viability was measured by MTT assay. SSMD was calculated relative to vehicle (DMSO) controls, with negative values indicating reduced viability. The red dashed line indicates the hit threshold (SSMD < −3.0). A total of 114 compounds met this criterion at one or both concentrations, indicating significant reductions in viability in both *AXL*^WT^ and *AXL*^KO^ cells relative to vehicle treated controls. Of these, 27 preferentially reduced viability in *AXL*^KO^ cells compared with *AXL*^WT^ cells. Compounds are listed in reverse alphabetical order, with each bar representing an individual compound at the indicated concentration. Compounds appearing under multiple library entries (e.g., dasatinib, dasatinib monohydrate, and dasatinib hydrochloride; AT7519 and AT7519 HCl; flavopiridol and flavopiridol HCl) were screened as independent entries and are displayed separately.

**Supplementary Figure 4: Classification of cancer cell lines by *AXL* expression for PRISM drug sensitivity analysis. (A)** Pathway-level enrichment of *AXL*^KO^ selective hits in the kinase inhibitor screen. The x-axis indicates the percentage of compounds within each annotated pathway that were selective hits (SSMD < −3.0 in *AXL*^KO^ but not *AXL*^WT^ cells). Point size represents the number of selective hits per pathway. The dashed line indicates the screen-wide selective hit rate (1.5%). Pathway enrichment was assessed using a one-sided hypergeometric test with Benjamini-Hochberg correction. ***FDR < 0.001. **(B)** *AXL* mRNA expression across cancer cell lines from the DepMap Public 24Q4 release is shown as batch-corrected log_2_(TPM + 1). Each point represents one cell line, and the box indicates the median and interquartile range. Cell lines were stratified into four groups according to *AXL* expression quartiles: *AXL*-Low (first quartile; log_2_(TPM + 1) < 0.4), *AXL*-Low-Med (first quartile to median; approximately 0.4-3.5), *AXL*-Med-High (median to third quartile; approximately 3.5-6.4), and *AXL*-High (above the third quartile; log_2_(TPM + 1) >6.4). *AXL*-Low (n = 40) and *AXL*-High (n = 155) cell lines included in the PRISM Repurposing Screen (20Q2) were selected for comparative drug-sensitivity analysis.

**Supplementary Figure 5: Dose-response validation of candidate inhibitors in *AXL*^WT^ and *AXL*^KO^ cells. (A)** Dose-response curves for 12 compounds in *AXL*^WT^ (grey) and *AXL*^KO^ (red) PC3 cells following 72 h of treatment. Cell viability was measured by MTT assay and normalized to vehicle-treated controls. Points represent mean ±SEM from 3-4 biological replicates. Two-parameter variable-slope model (Top = 100%, Bottom = 0%) to estimate absolute IC_50_ values; LogIC_50_ and Hill slope were estimated from the data. **(B)** IC_50_ values with 95% confidence intervals for *AXL*^WT^ (dark grey) and *AXL*^KO^ (orange) cells, ordered by increasing IC_50_ in *AXL*^KO^ cells. Open right-pointing markers indicate that nilotinib IC_50_ exceeded 5,000 nM in both cell lines. **(C)** AXL dependent sensitization, expressed as the *AXL*^WT^/*AXL*^KO^ IC_50_ ratio with 95% confidence intervals. Ratios >1 indicate greater sensitivity in *AXL*^KO^ cells. The dashed line indicates no shift (1.0-fold). Orange indicates significant shifts after Benjamini-Hochberg correction (adjusted *P* < 0.05); grey indicates non-significant shifts. Nilotinib was excluded because its fold shift could not be determined. Both x-axes are logarithmic. Fold-shift confidence intervals and P-values were derived from differences in log_10_(IC_50_) using a two-sided Wald test.

