## Supplemental Figures for "A Multi-Kinase Inhibitor Screen Identifies mTOR Signaling as a Vulnerability in Advanced Prostate Cancer"

Supplemental Figure 1

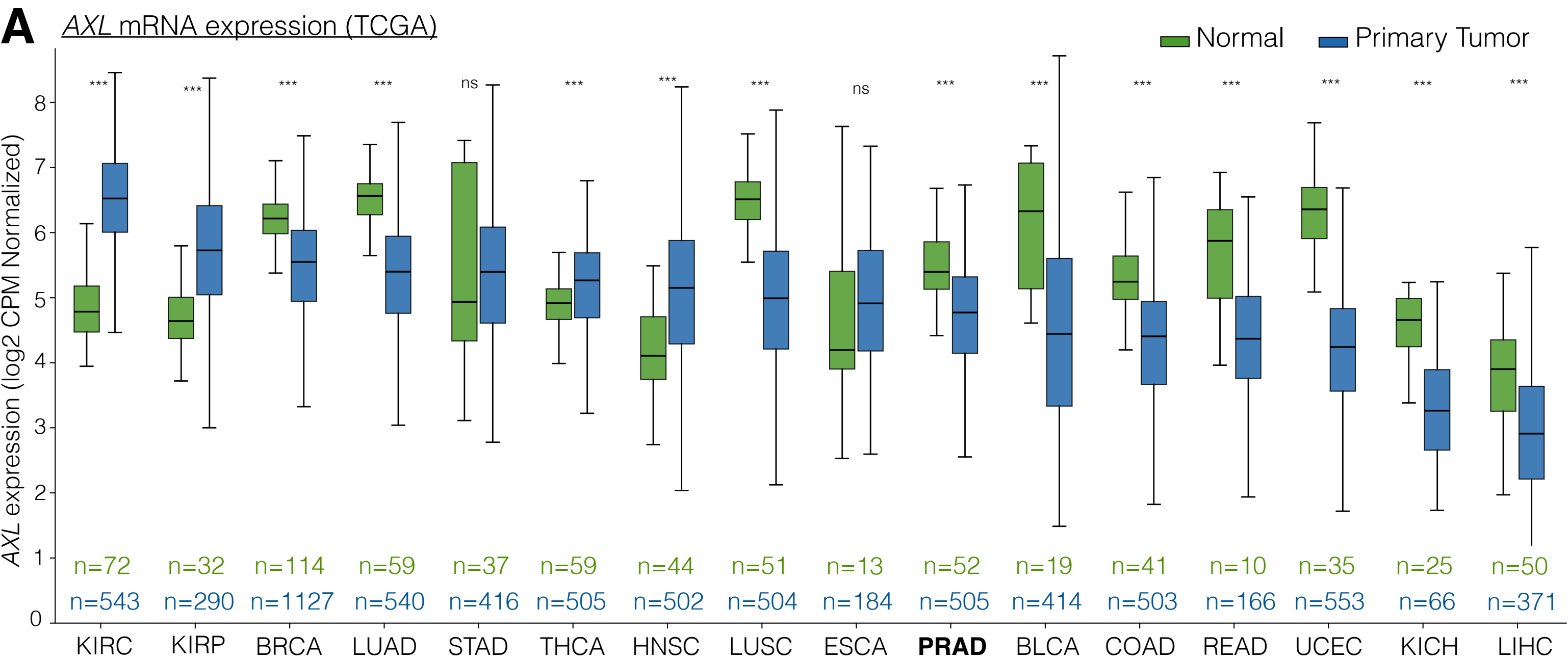

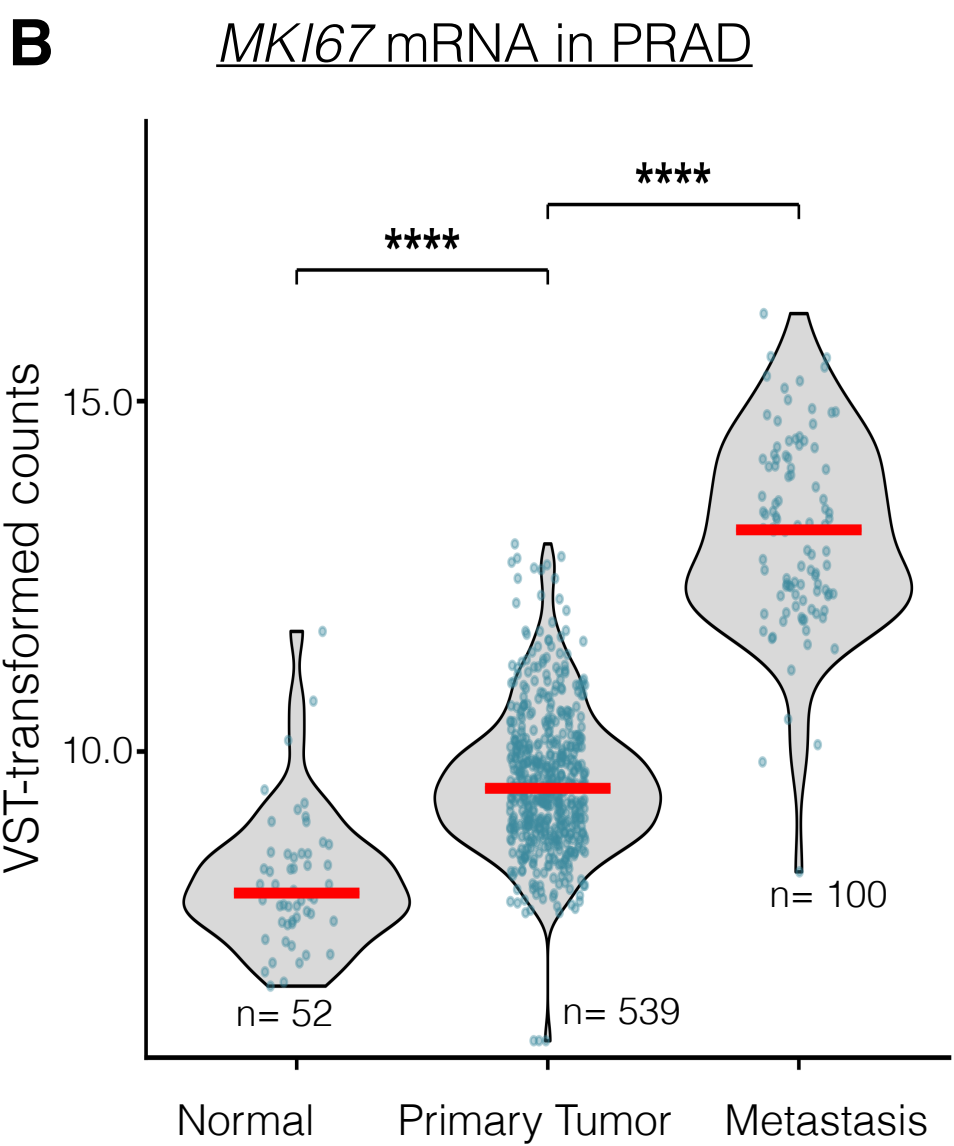

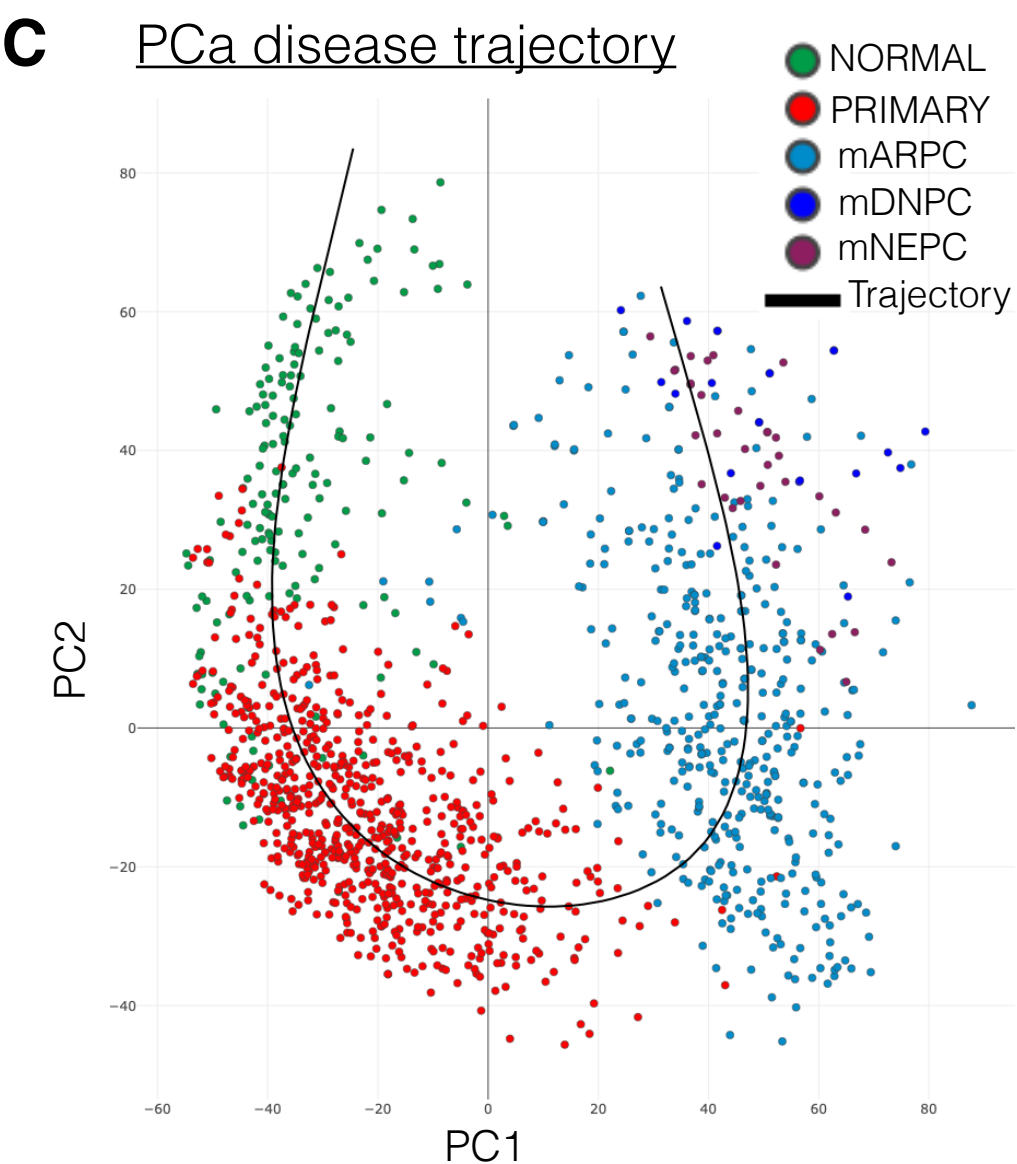

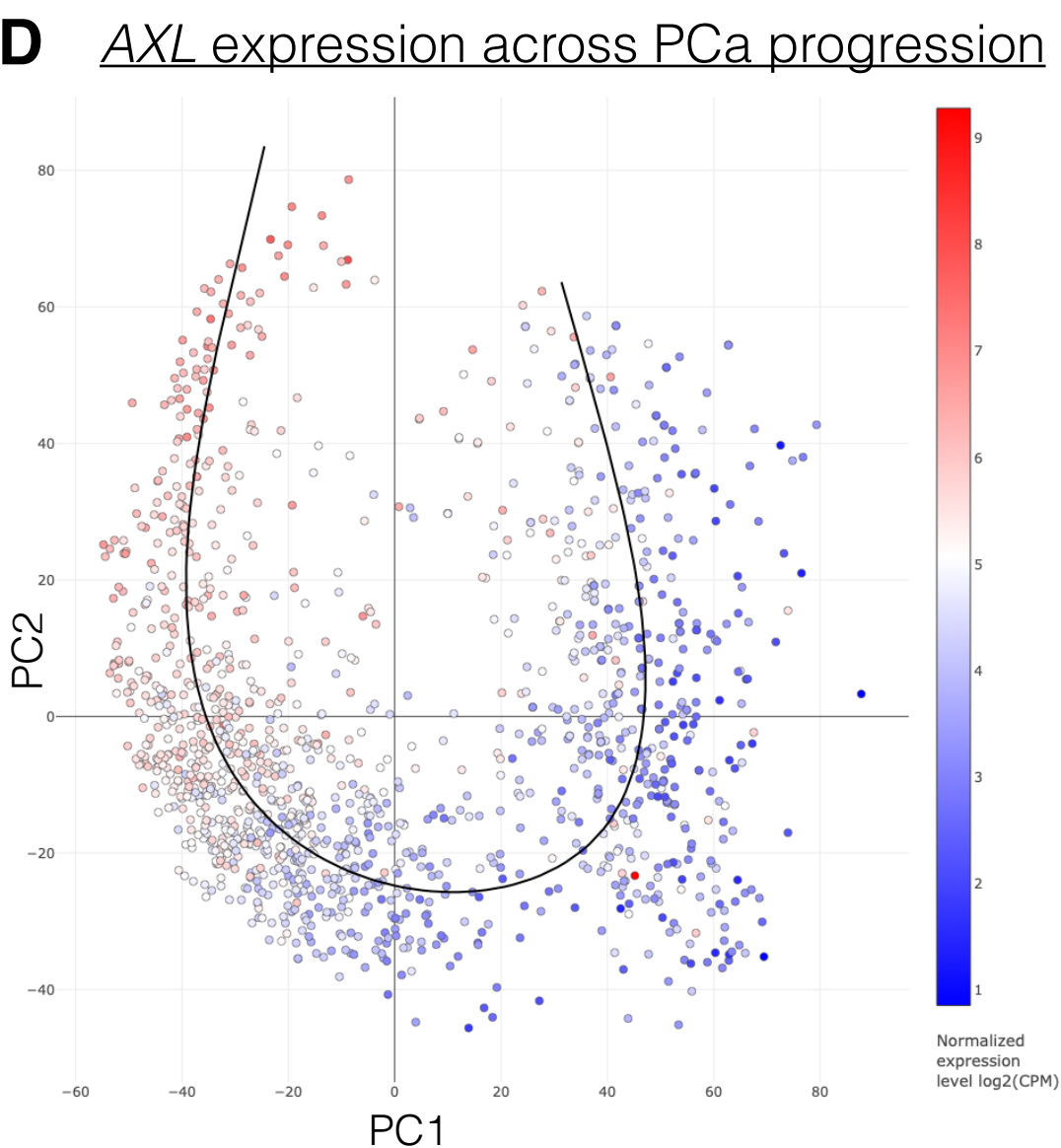

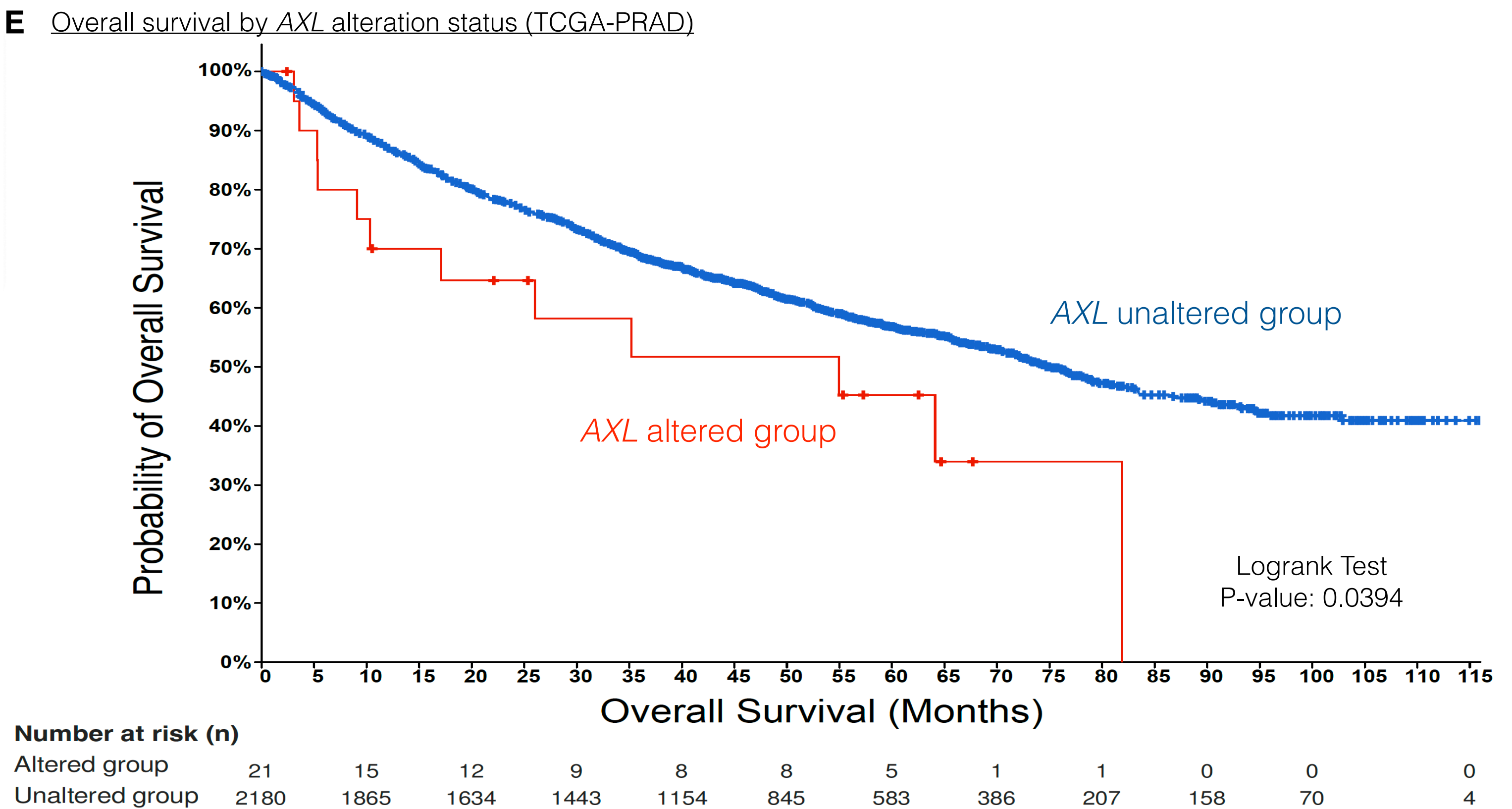

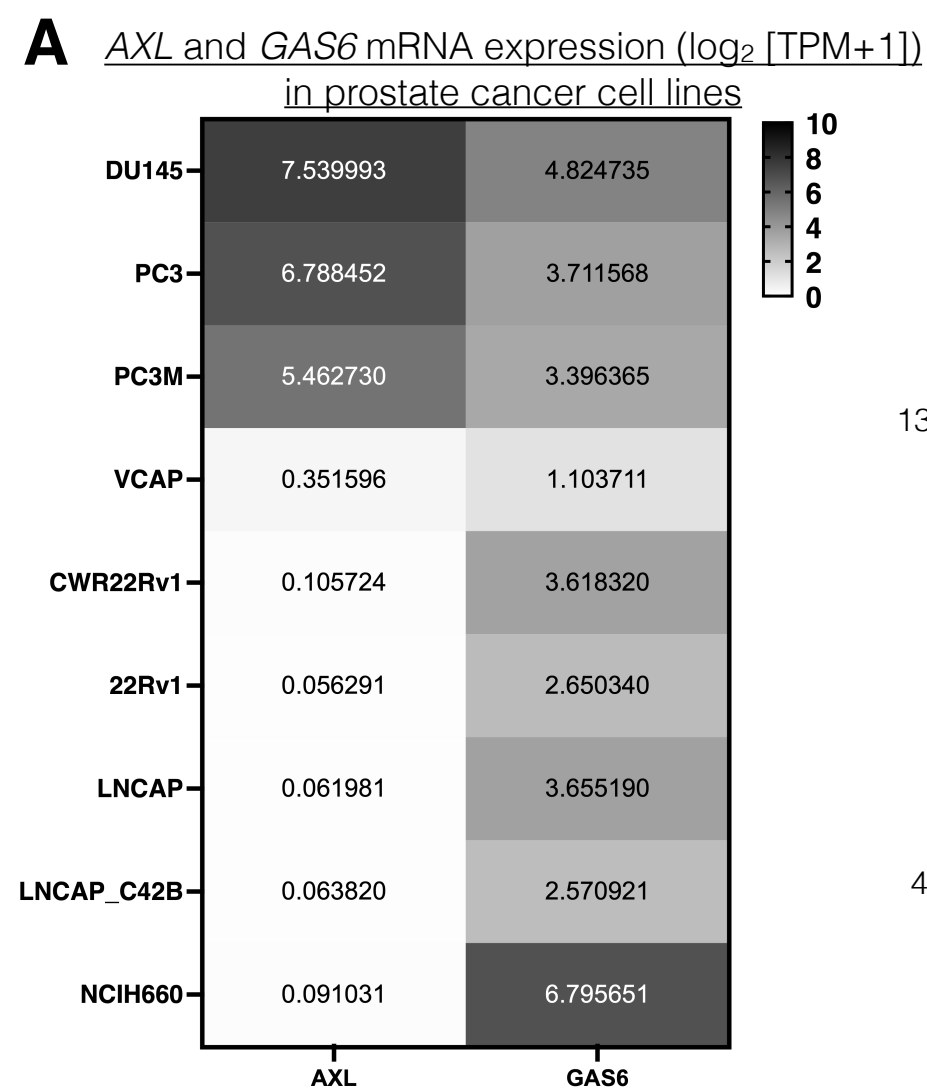

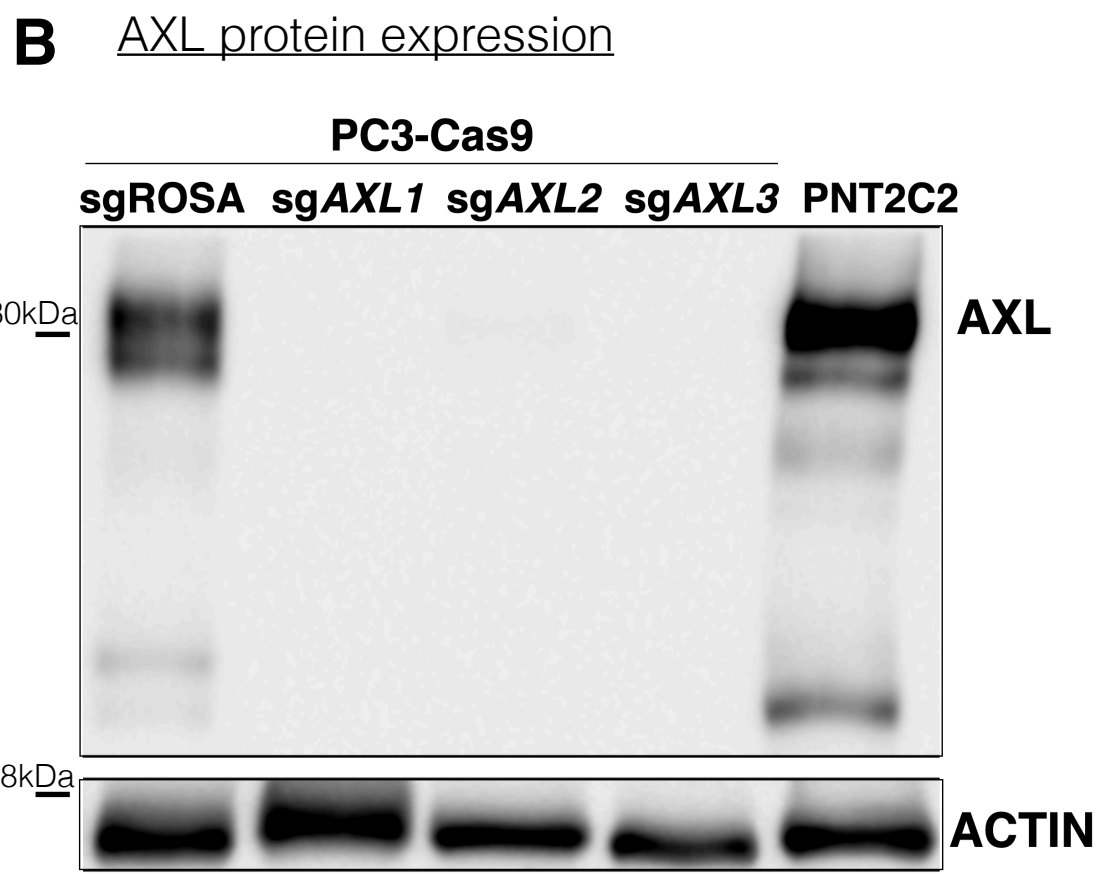

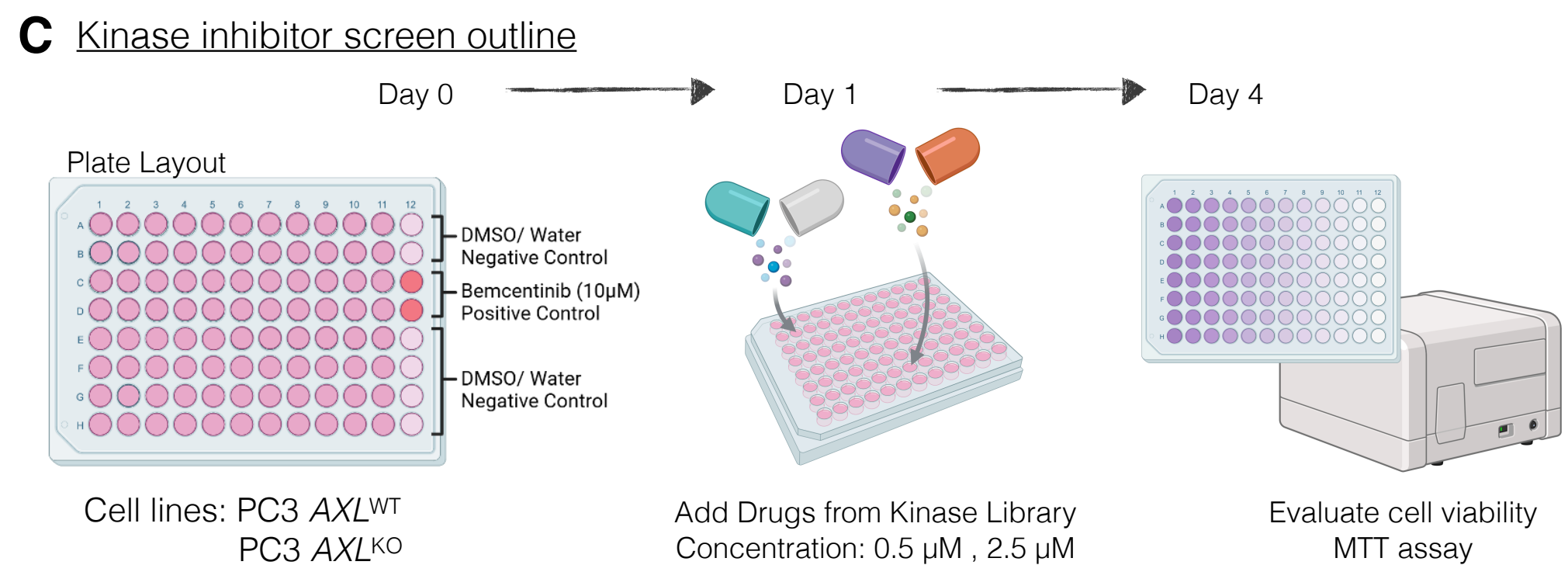

**D** Hit Identification using SSMD analysis (strictly standardized mean difference)

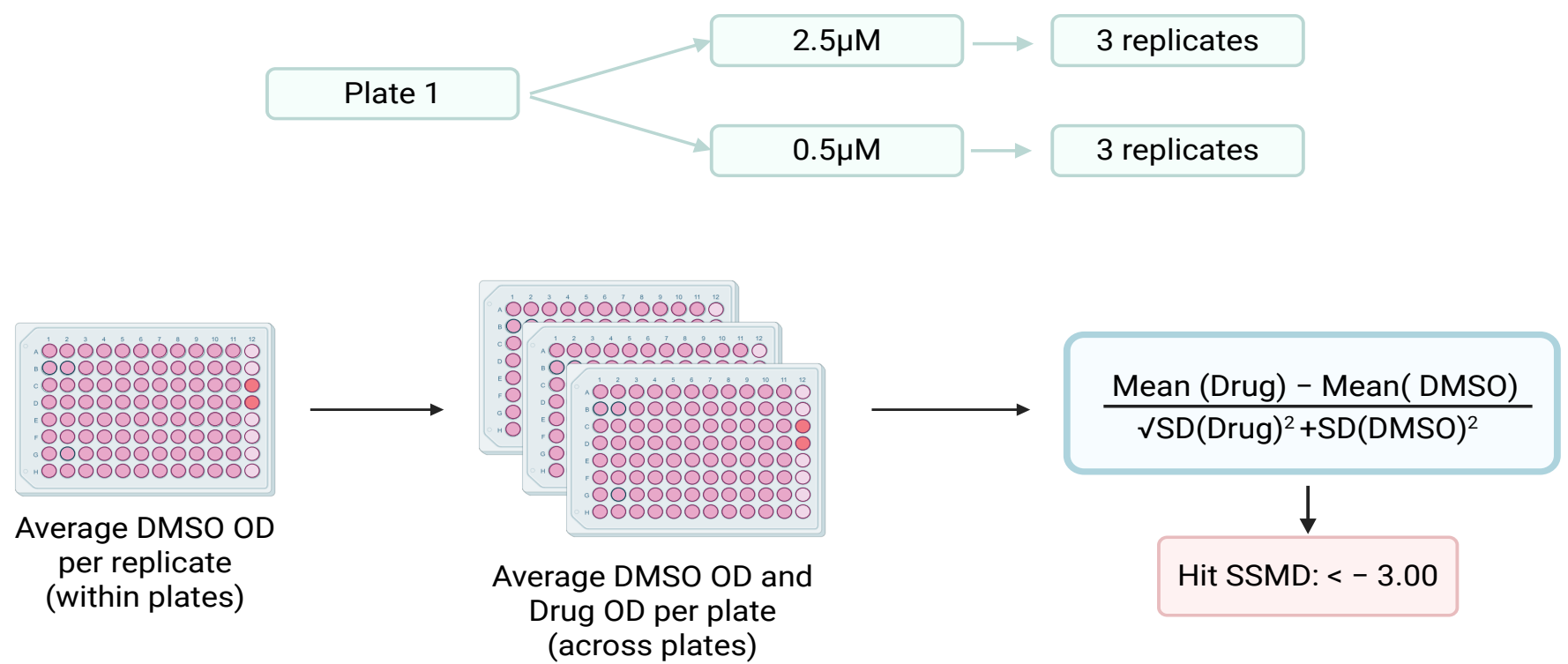

Supplemental Figure 3

Drugs

SSMD by drug

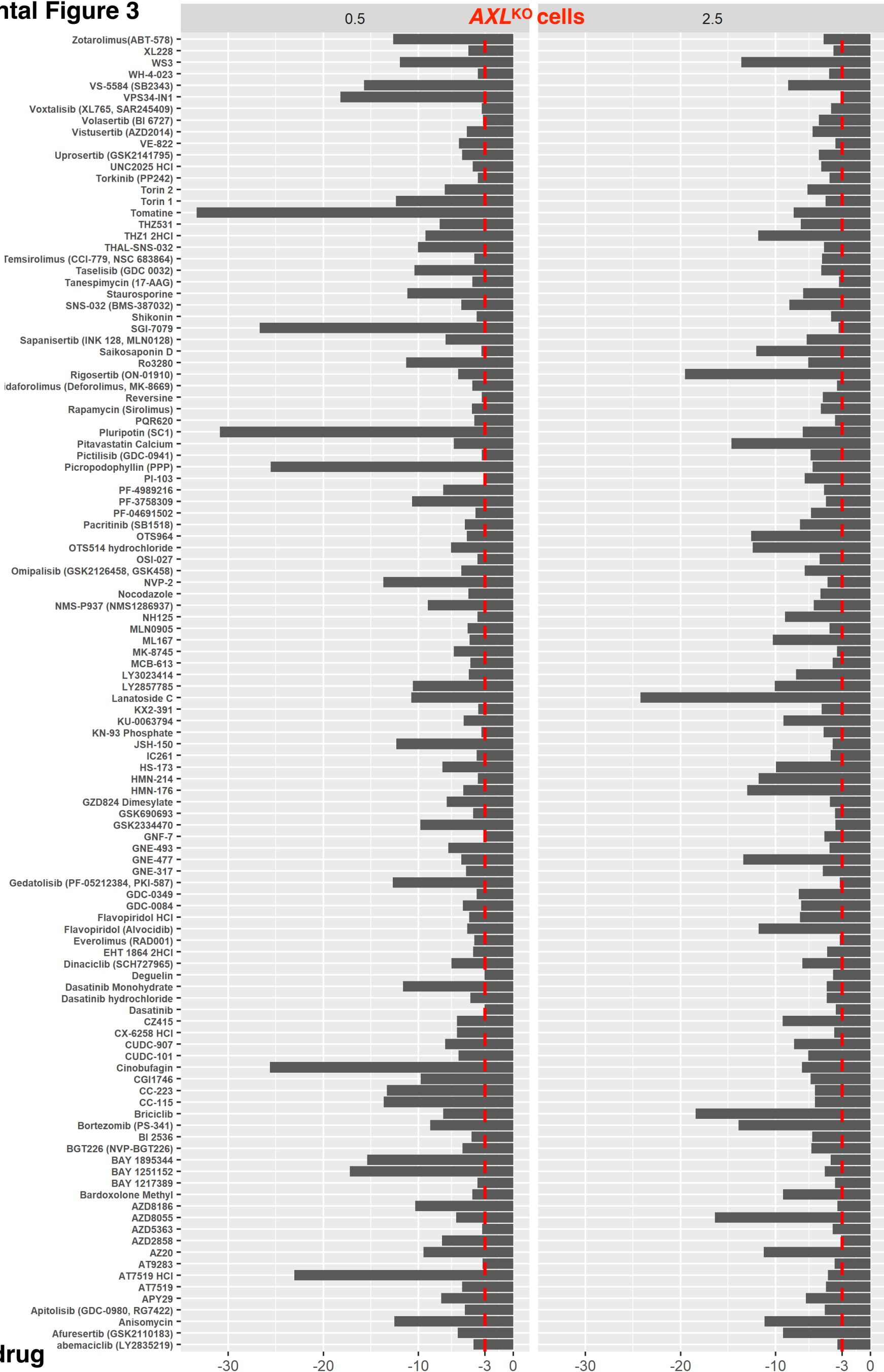

**A** *AXL*<sup>KO</sup> selective target pathway

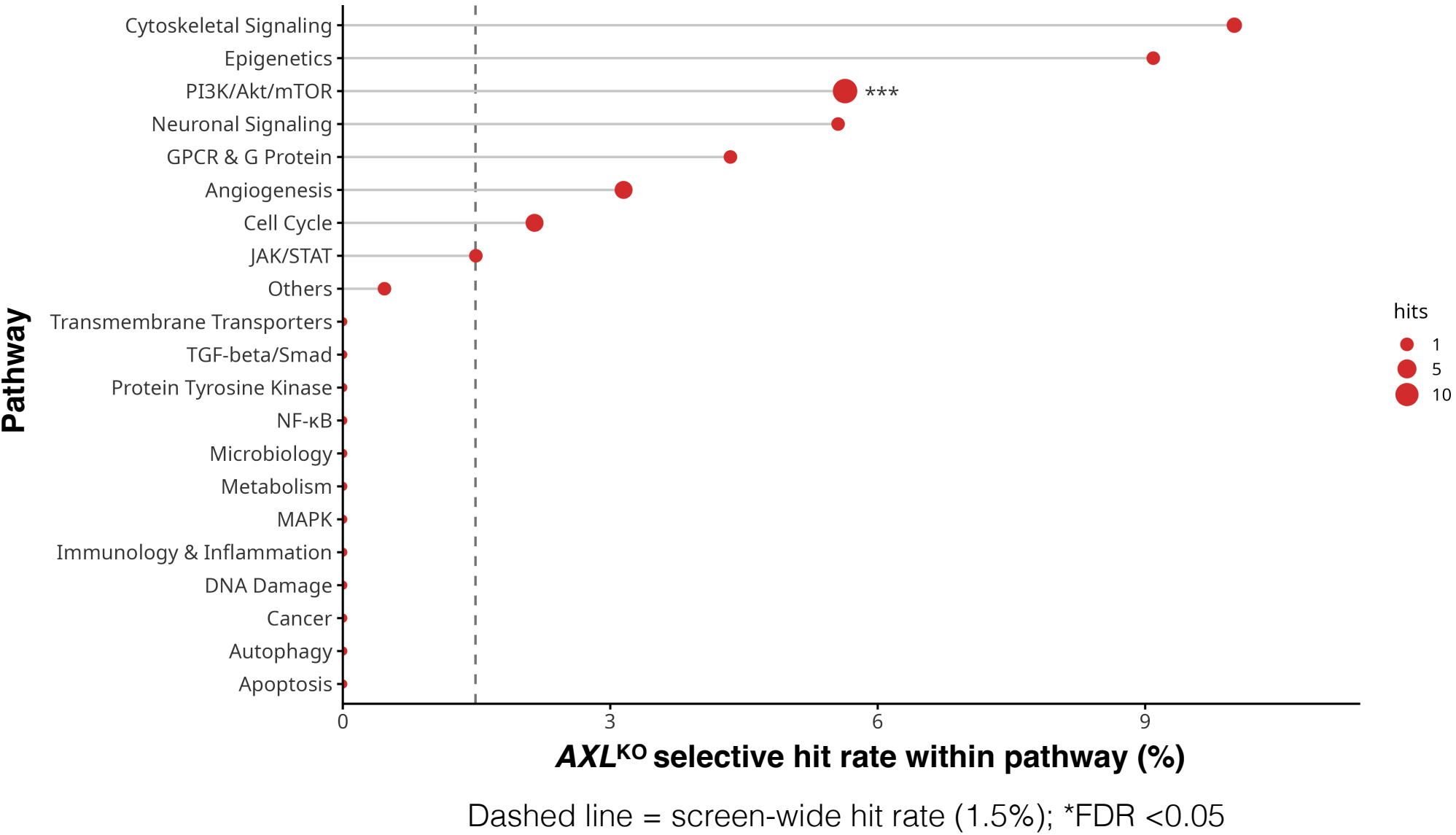

**B** DEPMAP: log2TPM *AXL* mRNA expression across cell lines

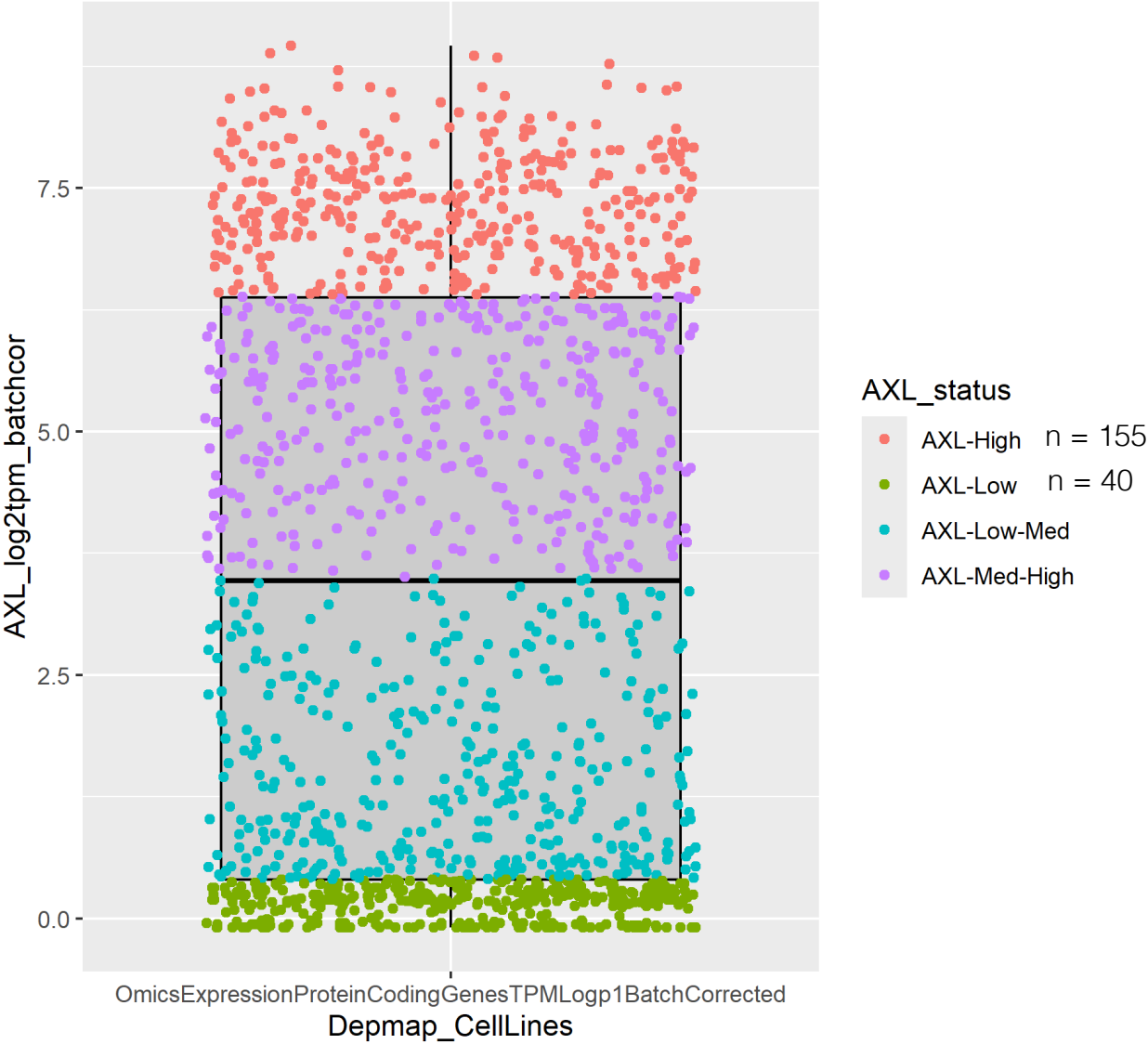

### Supplemental Figure 5

#### A Dose-response curves of candidate inhibitors

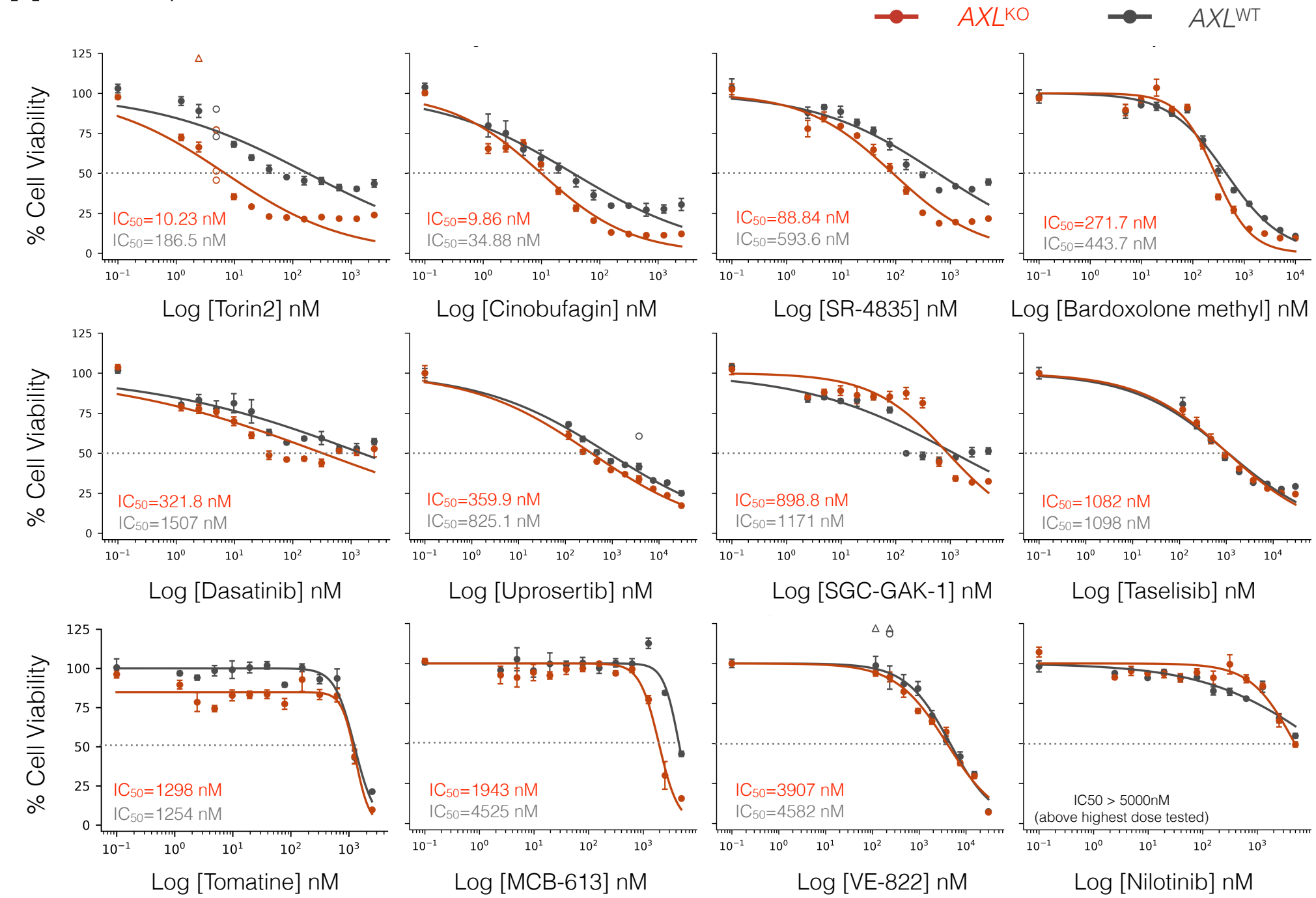

#### B Absolute potency

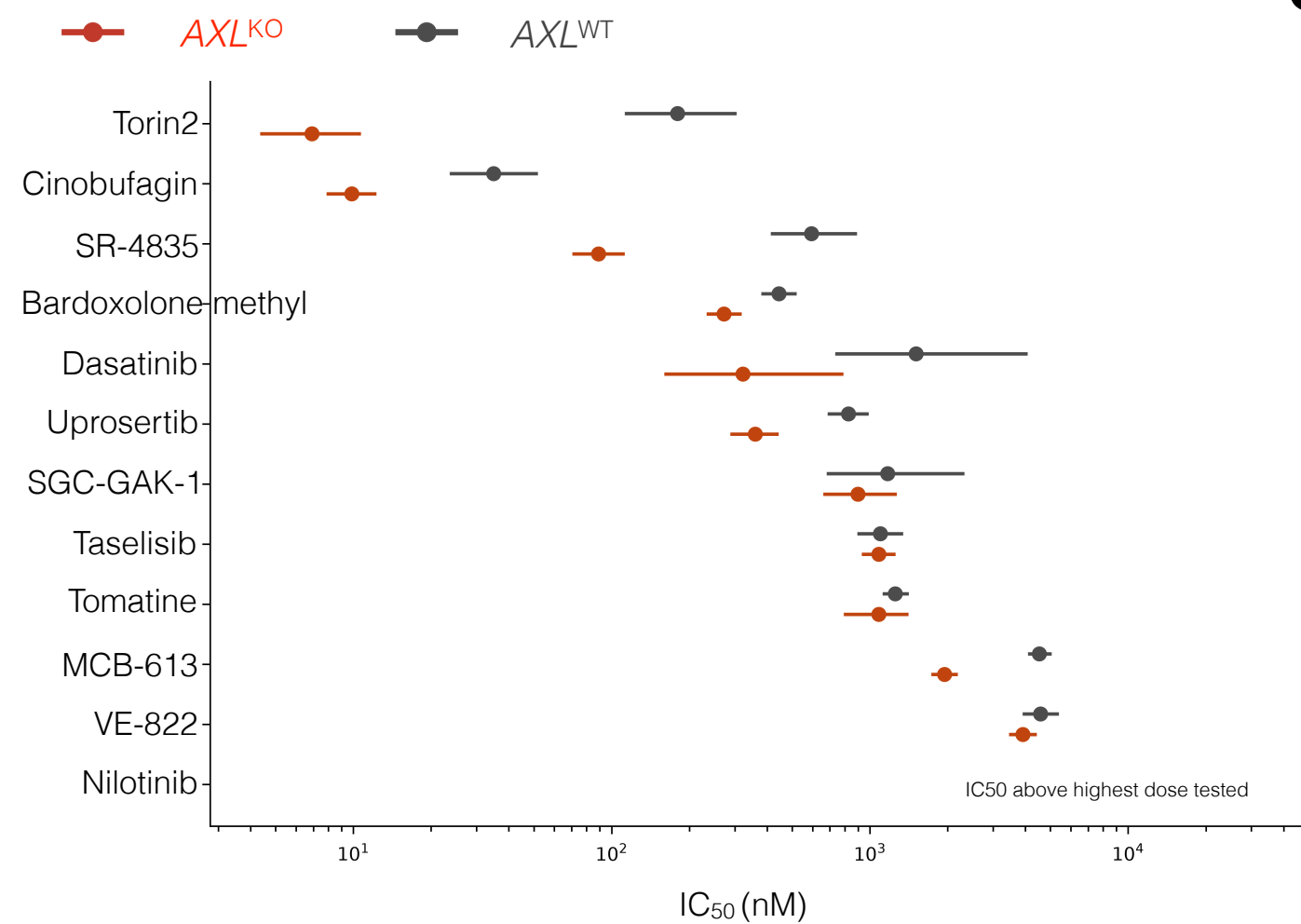

#### C AXL dependent Sensitization

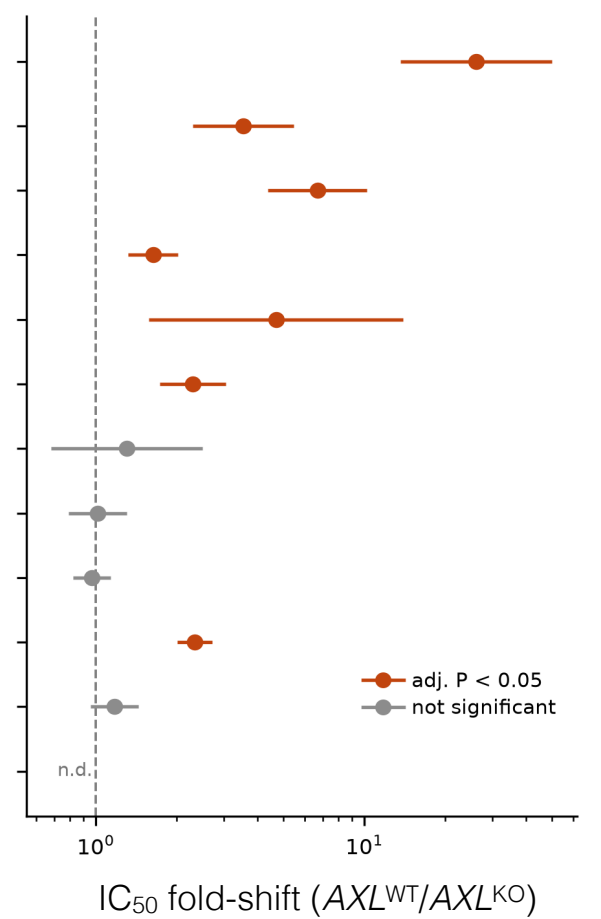
